# Aβ-HMGB1 complex is a pathogenic molecule at the advanced stage of Alzheimer’s disease

**DOI:** 10.1101/2025.06.12.659430

**Authors:** Hikari Tanaka, Hidenori Homma, Yuki Yoshioka, Kyota Fujita, Shingo Yamada, Katsutoshi Furukawa, Aiko Ishiki, Naoki Atsuta, Masaaki Waragai, Tatsuo Mano, Masahisa Katsuno, Gen Sobue, Hiroyuki Arai, Atsushi Iwata, Hitoshi Okazawa

**Author notes:** Contributed equally (Hikari Tanaka, Hidenori Homma, Yuki Yoshioka, Kyota Fujita, Shingo Yamada).

## Abstract

Multiple molecules including Aβ, tau and other inflammatory molecules mediate Alzheimer’s disease (AD) pathology. High mobility group box 1 (HMGB1), which is released from necrotic cells and binds to Toll-like receptors (TLRs) of surrounding neurons and microglia, also mediates AD pathology from the early stage. Paradoxically, HMGB1 concentration in cerebrospinal fluid (CSF) at the advanced stage of AD is not higher than that at the early stage. Here we show that Aβ-HMGB1 complexes are generated in neurons undergoing secondary necrosis around Aβ plaques in the AD brain at the advanced stage. Aβ-HMGB1 complex triggers neurite degeneration and necrosis of human normal iPSC-derived neurons via binding to TLR4. Further, two anti-HMGB1 antibodies that inhibit its interaction with TLR4 successfully suppress the toxicity of Aβ-HMGB1 complex to human iPSC-derived neurons, and recover cognitive impairment and Aβ-HMGB1 complex-related brain pathology in AD model mice, while such therapeutic effects were not obvious with an anti-Aβ antibody (lecanemab) approved for human AD patients. Enzyme-Linked Immuno Sorbent Assay (ELISA) revealed plasma level of Aβ-HMGB1 complex was increased in a part of AD patients at the advanced stage. These findings for the molecular basis of toxicity to neurons suggest the significance of Aβ-HMGB1 complex at the advanced stage of AD pathology, and might explain the discrepancy between Aβ burdens and clinical symptoms of human AD patients treated with anti-Aβ antibody.

## Main

HMGB1 protein, a DNA architectural protein that possesses two DNA binding domain called HMG box and a positively and negatively charged C-terminal regions, is located dominantly in the nucleus, while it shuttles between nucleus and cytosol^1,2^. Nuclear HMGB1 binds to minor grooves of DNA double strand and inserts charged tail between DNA and chromatin to unwind DNA double strand from chromatin for RNA transcription, DNA duplication, DNA damage repair and so on^1,2^. HMGB1 in cytosol contributes to autophagy^3^, mitochondrial quality control or mitochondrial DNA damage repair^4^. In addition to the intracellular roles, extracellular HMGB1 is a representative molecule for damage-associated molecular patterns (DAMPs)^5–7^ and a possible member of senescence-associated secretory phenotypes (SASPs)^8–11^.

After release to extracellular space, HMGB1 exists in three forms, disulfide (dsHMGB1), fully reduced/all-thiol (atHMGB1), and fully oxidized (oxHMGB1). Among them, receptors and biological functions of dsHMGB1 and atHMGB1 are well-characterized^12–14^. Especially, dsHMGB1 induces cell signaling of innate immune cells that finally leads to inflammation^5,6,14^ through binding with toll-like receptors (TLR)^15,16^ or receptor for advanced glycosylation end products (RAGE)^17^ Contrastingly, atHMGB1 forms a complex with C-X-C motif chemokine ligand 12 (CXCL12)^18^ to induce regeneration by binding to CXCR4^18–20^. Moreover, any type of HMGB1 could form heterogeneous complexes with multiple types of molecules, including ssDNA, LPS, IL-1, and nucleosomal proteins^21^.

Amyloid beta (Aβ) fragment is cleaved from amyloid precursor protein (APP) by β-secretase (Beta-site APP cleaving enzyme 1; BACE1)^22,23^ and the γ-secretase complex^24–26^ within the cell membrane and intracellular membranes of organelles such as the endoplasmic reticulum (ER) and Golgi apparatus^27^. HMGB1 secretion from the nucleus to the extracellular space is considered to be independent of the ER-Golgi system, as HMGB1 lacks the ER trafficking signal sequence, and the operative mechanism for this phenomenon remains unknown^28,29^. However, HMGB1 and Aβ could co-localize in the ER-Golgi system when the ER membrane becomes unstable in neurons undergoing necrotic processes such as transcriptional repression-induced atypical cell death (TRIAD), which is closely linked to the pathology of AD^30,31^, FTLD^32^, and senescence^31^. Recently, a morphological change highly similar to TRIAD named as PANTHOS was reported^33^, both of which occur at the ultra-early stage before extracellular Aβ deposition^30,33^. Moreover, another group reported that axonal spheroid full of LAMP1-positive vesicles formed by the autophagy-lysosomal degradation pathway underlies the early-stage cognitive dysfunction of AD patients and named the pathological change as PAAS^34^, whose morphology also seems similar to TRIAD and PANTHOS.

The roles of innate immunity and microglia in AD pathology have become a significant recent research interest since a large portion of genetic risk factors of AD turned out to be concentrated at innate immunity functions ^35^ Disease-associated microglia (DAM) that inhibit propagation of Aβ and potentially Tau^36^ and are regulated by TREM2, a genetic risk factor for AD^27,37^, is a good example supporting the significance of microglia^38^. However, microglia in the brain are heterogeneous^36,39,40^ and some subpopulations enhance brain inflammation by PQBP1-mediated recognition of Tau as a pathogen and activating the cGAS-STING-NFκB pathway^41,42^. In addition to Tau and Aβ, HMGB1 released from necrotic cells critically impacts microglia through TLRs^5,16,43,44^.

HMGB1 also binds TLRs on the surfaces of neurons, and induces degeneration of the neurite and cell body via phosphorylation of MARCKS and Ku70, respectively^30,45^. A number of reports have also shown that soluble Aβ such as oligomers or protofibrils degenerate synapse on neurites^46–50^. Interestingly, Aβ and HMGB1 form various heterogenous complexes^45^. Though such Aβ-HMGB1 complexes might play a certain role, the pathological effect on microglia and neurons, the clinical relevance to progression of AD pathology as well as the generation site in brain of the Aβ-HMGB1 complex have not been elucidated.

In this study, we address the role of Aβ-HMGB1 complex. Aβ-HMGB1 complexes are formed mainly in the ER-Golgi system of cortical neurons in AD pathology and that Aβ-HMGB1 complex triggers neurite and neuronal degeneration like HMGB1. Aβ-HMGB1 complex is detectable in blood and cerebrospinal fluid (CSF) samples of human AD patients. Aβ-HMGB1 complex acts directly on neurons to induce neurodegeneration via binding to the pattern recognition receptor Toll-like receptor 4 (TLR4) similarly to HMGB1^30,31,45,51^, and the effect is independent of the actions of Aβ in both the fibril^52^ and dimer states^53^. Further, anti-HMGB1 antibody shown effective on AD mouse models^31,45^ but not the anti-Aβ-antibody lecanemab inhibits neurodegeneration of human iPSC-derived neurons induced by Aβ-HMGB1 complex.

### Aβ-HMGB1 complexes are formed *in vitro* and present in AD brains *in vivo*

First, we reexamined whether Aβ and HMGB1 complexed *in vitro*. Pre-incubation of Aβ and HMGB1 at a 1:1 molar ratio at 37□ for 48 hours *in vitro* followed by Western blot with anti-Aβ and anti-HMGB1 antibodies confirmed the presence of an Aβ-HMGB1 complex of approximately 35 kDa, which is the predicted molecular weight of Aβ:HMGB1 = 1:1 (Fig. 1a, red arrow). Heterodimers were detected in mixtures of HMGB1 with either Aβ1-40 or Aβ1-42 (Fig. 1a), while formation of the heterodimer was less robust in the mixture of Aβ1-42 with HMGB1 (Fig. 1a). Reaction of the 6E10 antibody with the Aβ monomer was much stronger than that of the 82E1 antibody (Fig. 1a), while both antibodies detected the heterodimer. More oligomers and protofibrils were synthesized with Aβ1-42 than with Aβ1-40, and their synthesis was inhibited by the presence of HMGB1 (Fig. 1a, blue arrowheads).

**Figure 1.**
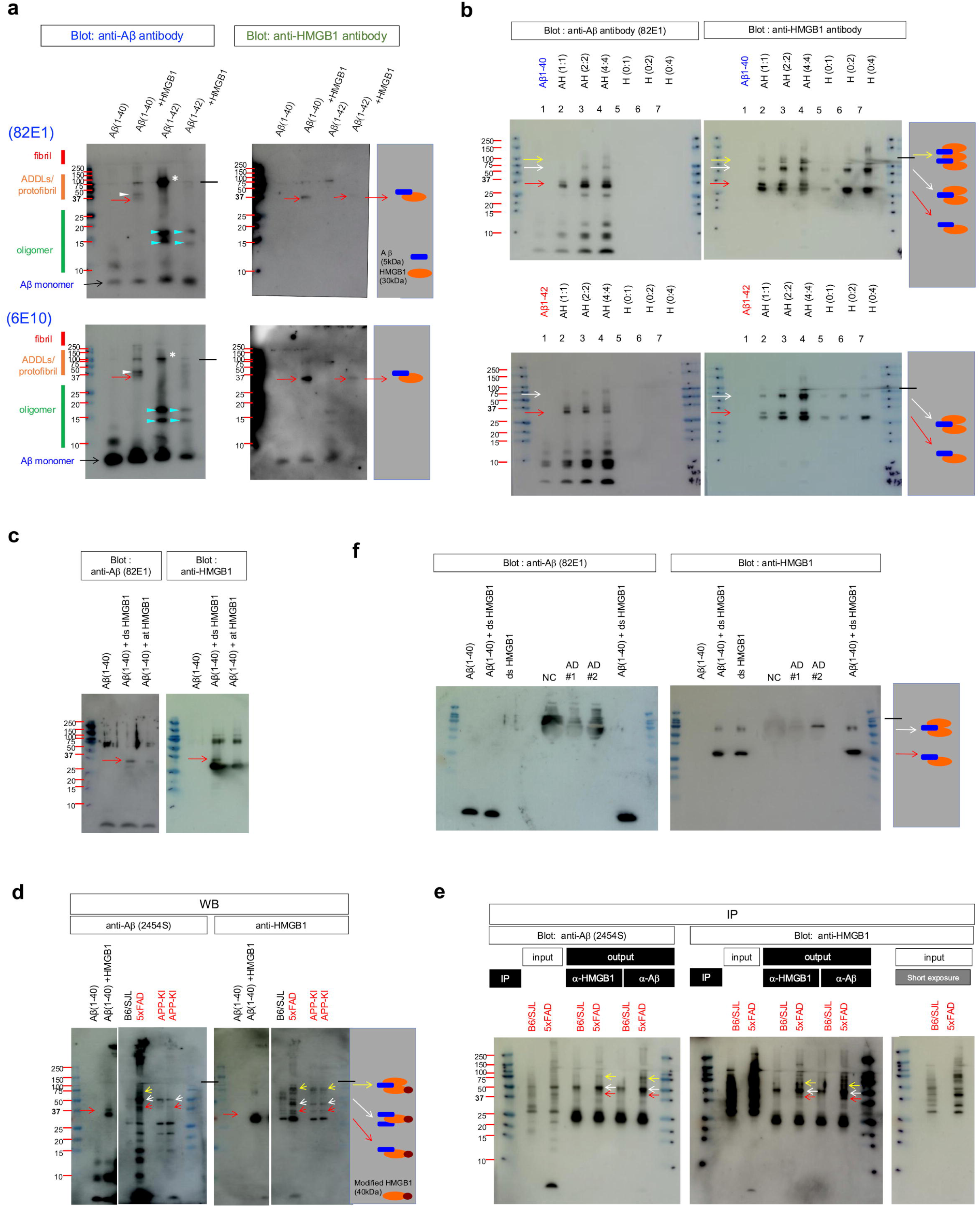
*In vitro* and *in vivo* presence of Aβ-HMGB1 complex. a) Aβ1-40 peptide or Aβ1-42 peptide was incubated with the mixture of recombinant human HMGB1 proteins (R&D, 1690-HMB-050), including dsHMGB1 and atHMGB1, for 48 hours at 37°C, and the synthesized protein was detected by Western blot with anti-Aβ antibody (82E1 or 6E10) and subsequently with anti-HMGB1 antibody. A heterodimer (red arrow) was detected. More oligomers (blue arrowheads) and protofibrils (white asterisk) were formed with Aβ1-42 than with Aβ1-40, while Aβ-HMGB1 complex synthesis was inhibited in the presence of HMGB1. The band above Aβ-HMGB1 heterodimer (white arrowhead) might be a heterotrimer (two Aβ + one HMGB1). b) Aβ1-40 or Aβ1-42 was mixed with dsHMGB1 (HMGBiotech, HM-120) at the specified concentrations and incubated for 24 hours at 37°C. The mixtures of Aβ-HMGB1 complexes were separated by SDS-PAGE with a gradient gel, and blotted with anti-HMGB1 antibody and subsequently with anti-Aβ antibody (82E1). The border line of the gradient gel is indicated with a black line. The heterogeneous dimer of Aβ-HMGB1 complex is indicated with a red arrow, the trimer is indicated with a white arrow, and the pentamer is indicated with a yellow arrow. c) Aβ-HMGB1 complex generation with dsHMGB1 or atHMGB1 was compared. The separate forms of HMGB1 were incubated with Aβ1-40 for 48 hours and subjected to SDS-PAGE. The same filter was forwarded to Western blot with anti-HMGB1 antibody and subsequently with anti-Aβ antibody (82E1). A greater amount of Aβ-HMGB1 complex was generated with dsHMGB1 than with atHMGB1 under these conditions. d) Immunoblots of total cerebral cortex lysates of AD mouse models. Blotting with anti-Aβ antibody (2454S) and anti-HMGB1 antibody revealed the presence of multiple forms of Aβ-HMGB1 complexes *in vivo*. Western blot analysis revealed that 45 kDa (red arrow), 50 kDa (white arrow), and 75 kDa (yellow arrow) bands were increased in total cerebral cortex lysates of 5xFAD and APP-KI mice relative to control mice (B6/SJL), and the complexes were reactive to both anti-Aβ and anti-HMGB1 antibodies. nti-HMGB1 antibody revealed co-precipitation of HMGB1 and Aβ at 45 kDa (red arrow), 50 kDa (white arrow), and 75 kDa (yellow arrow), which corresponded to the bands detected in Western blot (Fig. 1d).

Aβ-HMGB1 complexes at higher molecular weights were also detected when the concentrations of Aβ and HMGB1 were increased in a 24-hour incubation (Fig. 1b). Both Aβ1-42 and Aβ1-40 formed a 65 kDa complex with HMGB1 (Fig. 1b, white arrow), which is equivalent to the molecular weight of Aβ: HMGB1 = 2:1. Aβ1-40, but not Aβ1-42, formed a third complex with HMGB1 at 100 kDa, which could be a hetero-pentamer composed of three HMGB1 molecules and two Aβ molecules (Fig. 1b, yellow arrow). Co-incubation of atHMGB1 with Aβ1-40 for 48 hours did not generate the 35 kDa complex as robustly as dsHMGB1 (Fig. 1c).

Subsequently, we determined whether the Aβ-HMGB1 complex was present in AD model mouse brains (Fig. 1d, e). Western blot analysis revealed 45 kDa, 50 kDa, and 75 kDa bands that were clearly increased in the cerebral cortex of 5xFAD and APP-KI mice relative to control B6/SJL mice (Fig. 1d). Immunoprecipitation of cerebral cortex lysates from AD mice with anti-Aβ or anti-HMGB1 antibody revealed that these bands were reactive to anti-HMGB1 and anti-Aβ antibodies, indicating that they represented Aβ-HMGB1 complexes (Fig. 1e). HMGB1 undergoes multiple modifications such as acetylation^54^, phosphorylation^55,56^, and methylation^4^. Interestingly, 2D electrophoresis analysis of HMGB1 purified from the thymus revealed 30 kDa, 35 kDa and 60 kDa HMGB1 protein spots^54^, suggesting that the 45 kDa, 50 kDa and 75 kDa bands detected in the present study could be complexes of these modified HMGB1 proteins.

### Aβ-HMGB1 complex is present in cells surrounding Aβ plaques

To investigate the pathological significance of the increase of Aβ-HMGB1 complex, we performed immunohistochemistry of two AD mouse models, 5xFAD (Supplementary Fig 1a, Supplementary Video 1) and APP-KI (*App*^NL-G-F/NL-G-F^-KI) mice (Supplementary Fig 1b, Supplementary Video 2), both at 6 months old. We previously classified phenotypes of cell death pathology into three patterns: active necrosis (primary necrosis of a single neuron), secondary necrosis, and ghost of cell death^30^. In both 5xFAD and APP-KI mice, co-localization of Aβ and HMGB1 signals was observed in neurons undergoing secondary necrosis at the periphery of extracellular Aβ aggregates (Aβ plaques) (Supplementary Fig. 1 Supplementary Fig. 1). Merged signals were present in the cytoplasm of dying cells with a faint DAPI signal (Supplementary Fig. 1a, b, Supplementary Video 1, 2) suggesting that the Aβ-HMGB1 complex formed in neurons secondarily damaged around senile plaque formation, which were characterized previously^30^. Quantitative analyses of the number of cells with Aβ-HMGB1 co-staining in 5xFAD and APP-KI mice confirmed the phenotype specific to AD mouse models (Supplementary Fig 1a, b, lower graphs).

Next, we examined postmortem cerebral cortexes of familial AD patients carrying the M146L mutation in the presenilin-1 (PS1) gene. Co-localization of Aβ and HMGB1 was present in the cytoplasm of cells surrounding extracellular Aβ aggregates (senile plaques) together with faint DAPI signals (Supplementary Fig 1c). HMGB1 was located in the nuclei of normal cells without intracellular Aβ (Supplementary Fig 1c, white arrow), while HMGB1 shifted from the nucleus to the cytoplasm in abnormal cells and co-localized with intracellular Aβ (Supplementary Fig 1c, squared). These morphological changes of abnormal neurons corresponded with typical features of TRIAD necrosis linked to impaired YAP function identified in HD and AD pathologies^30,57–59^. Quantitative analyses of the number of cells with Aβ-HMGB1 co-staining in human postmortem AD brains confirmed phenotypes specific to AD pathology (Supplementary Fig 1c, right upper graph). Moreover, we examined postmortem brains of non-familial AD patients. We detected two patterns of co-localization of Aβ and HMGB1. Robust signals of co-localization were found adjust to the core of Aβ plaque (Supplementary Fig 1d, #1), while the cytoplasmic co-localization of Aβ and HMGB1 surrounded vacuolar changes, which was similar to the pattern detected in PS1-linked AD patients, was also found at a distance from Aβ plaque (Supplementary Fig 1d, #2). Obviously, the #1 pattern seemed microglia and the #2 pattern looked like neurons.

### Aβ-HMGB1 complex is generated in dying neurons and scavenged by microglia

To confirm the cell type responsible for generation of Aβ-HMGB1 complex, we performed immunostaining with co-staining for MAP2, Iba1, or GFAP (Supplementary Fig. 1g-j, Supplementary Fig. 2). First, immunostaining with the neuron-specific marker MAP2 indicated that Aβ and HMGB1 co-localized in dying neurons surrounding Aβ plaques of 5xFAD mice (Supplementary Fig. 1g) and of APP-KI mice (Supplementary Fig. 1h). Moreover, in human AD patients carrying the PS1 mutation (Supplementary Fig. 1i) or in sporadic AD patients (Supplementary Fig. 1j), we detected co-localization of Aβ and HMGB1 in a single dying neuron with intracellular Aβ (Supplementary Fig. 1i, j), which was consistent with primary necrosis defined previously^30^ and in dying neurons at the periphery or in the vicinity of senile plaques, which is consistent with secondary necrosis defined previously^30,31^.

Immunostaining with the microglia-specific marker Iba1 indicated that the Aβ-HMGB1 complex could also be present in a population of microglia localized to the border of extracellular Aβ aggregates in 5xFAD mice (Supplementary Fig. 2a), APP-KI mice (Supplementary Fig. 2b), and human AD patients (Supplementary Fig. 2c). However, considering that microglia themselves do not generate Aβ from cleavage of APP and that microglia function as phagocytes, it would be reasonable to conclude that these microglia are phagocytosing and degrading Aβ-HMGB1 complex generated by clusters of necrotic neurons that later form Aβ plaques^30,31^.

Immunostaining for the astrocyte-specific marker GFAP did not reveal definite co-staining of Aβ and HMGB1 in either AD mouse model (Supplementary Fig. 2d, e). However, in the parietal lobe cortexes of human PS1-linked AD patients, reactive astrocytes with enlarged cytoplasm and some astrocyte processes contained merged Aβ and HMGB1 signals, indicating that during the course of long-term AD pathology, reactive astrocytes became involved in the metabolism of Aβ-HMGB1 complex. Morphological features suggested that astrocytes incorporating Aβ-HMGB1 complex by phagocytosis were reactive and induced inflammation, similar to other reported case^60^.

To determine the predominant cell types with Aβ-HMGB1 co-staining, we calculated the percentages of Aβ-HMGB1 co-stained cells in neurons, microglia, and astrocytes among the total numbers of each cell type. The percentages of each subtype were very similar in 5xFAD mice, APP-KI mice, and human AD patients, with approximately 60–80% in neurons, 0–10 % in astrocytes, and 80% in microglia. Considering the function, microglial Aβ-HMGB1 complex would be the result of phagocytosis, and microglia would not release Aβ-HMGB1 complex to extracellular space.

Collectively these findings indicated that Aβ-HMGB1 complex was generated primarily in the cytoplasm of neurons accumulating intracellular amyloid and undergoing the process of TRIAD necrosis described previously, which is induced by the HMGB1-TLR4-PKC-Ku70 pathway^30,31^. Neurons undergoing TRIAD necrosis exponentially increase via the mechanism of “amplification from necrosis to necrosis” during the progression of AD pathology^30,31^. Because the Aβ burden also increases during the progression of AD pathology, Aβ-HMGB1 complex generation would likely increase at advanced AD stages, and localization of the Aβ-HMGB1 complex generation site to secondary TRIAD neurons would explain the high values of Aβ-HMGB1 complex in the advanced stages of AD but not in the MCI stage.

### Aβ-HMGB1 complex is generated in ER-Golgi system of neurons

We further addressed subcellular organelles where Aβ-HMGB1 complex was generated by immunohistochemistry of AD mouse models with various organelle markers. GM130, a marker for Golgi apparatus, consistently stained the intracellular regions both reactive for Aβ and HMGB1 (Fig. 2a). KDEL, a marker for ER, partially overlaid on the double positive area of Aβ and HMGB1 (Fig. 2a), whereas Aβ-HMGB1 merged signals were observed in the area adjacent to but deficient of KDEL stains in some cases, which might be Golgi apparatus (Fig. 2a). We also found that a part of LC3-positive areas were also stained with Aβ and HMGB1 (Fig. 2a). Consistently, we confirmed these Aβ-HMGB1 merged signals at ER, Golgi apparatus and autophagosome in human sporadic AD patients (Fig. 2a). In contrast, Aβ-HMGB1 complex signals did not merge with the mitochondrial marker protein ATP5A in cortical neurons of two mouse AD models and of human sporadic AD patients (Fig. 2b). These results suggested that Aβ-HMGB1 complex was generated mainly in ER-Golgi system that also plays an essential role in production of Aβ peptides cleaved from APP inserted in organelle membranes of the ER-Golgi system^61–63^.

**Figure 2.**
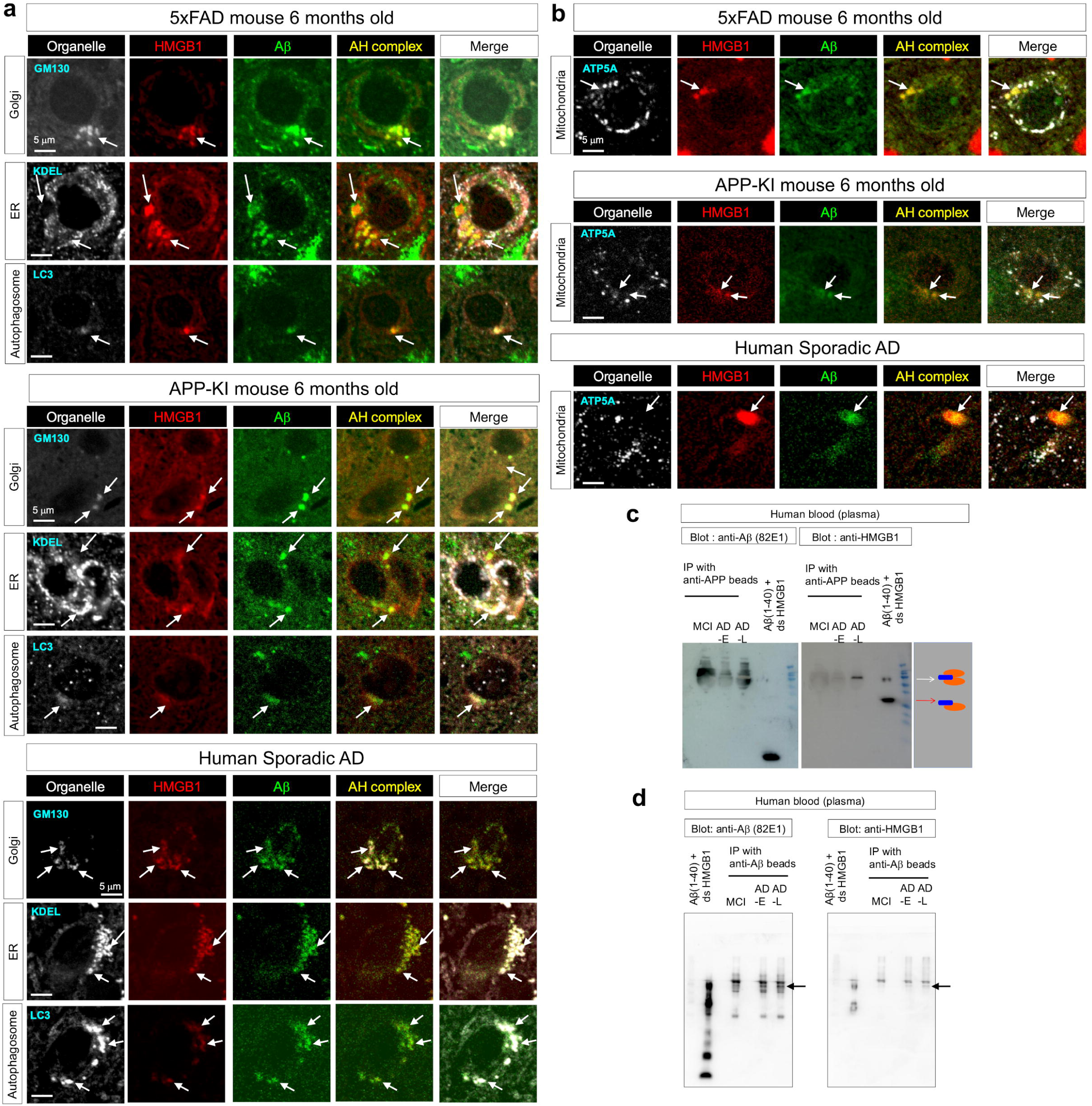
Colocalization of Aβ and HMGB1 in intracellular organelles. a) Organelle localization of Aβ-HMGB1 complex was analyzed by immunohistochemsitry using Golgi apparatus-specific (GM130), endoplasmic reticulum-specific (KDEL) and autophagosome-specific (LC3) markers in 5xFAD mice (upper panels) or APP-KI mice (middle panels). In both AD model mice, Aβ and HMGB1 were co-localized in Golgi apparatus and autophagosome but not in endoplasmic reticulum. Immunohistochemistry of cerebral cortex of sporadic cases AD patients. Organelle localization of Aβ-HMGB1 complex in human sporadic AD patients was similar to that in AD mouse models (lower panels). b) Immuohistochemistry of mouse AD models and human sporadic AD patients with a mitochondria-specific marker (ATP5A). c) Immunoprecipitation from plasma of MCI, sporadic early-stage AD, sporadic late-stage AD patients with anti-APP beads. The sample of revealed existence of HMGB1 in the APP complex. d) Immunoprecipitation from plasma of sporadic AD patients with anti-Aβ beads revealed existence of HMGB1 in the complex with Aβ.

Golgi apparatus releases intraluminal substances outside of cells by generating secretory vesicles or by providing intracellular vesicles that fuse with to endosomes to form multivesicular bodies (MVBs) and are finally released from cells as exosomes^64^. These considerations prompted us to test whether APP and HMGB1 co-exist in secreted vesicles in peripheral blood. Anti-APP N-terminal antibody-conjugated beads were employed instead of Aβ antibody to precipitate APP-containing vesicles specifically (Fig. 2c). The precipitation was performed without using detergent, and samples precipitated from plasma samples of MCI (ADAS-cog 8.6), early stage AD (MMSE 27) and advanced stage AD (MMSE 13) patients included various lengths of N-terminal peptides and their aggregates containing Aβ regions (Fig. 2c). Interestingly, we detected a band reacting with anti-HMGB1 antibody and corresponding to a type of Aβ and HMGB1 complex (two HMGB1 and one Aβ peptide), which was detected in a positive control generated by in vitro incubation, only in the sample from advanced-stage AD patient (Fig. 2c). Moreover, we precipitated the Aβ-HMGB1 complex by anti-Aβ-antibody-conjugated beads, and confirmed existence of the similar type of Aβ-HMGB1 complex in AD patient plasma by western blot of the precipitate (Fig. 2d).

### Aβ-dsHMGB1 complex induces neurite degeneration and neuronal death

We examined whether Aβ-HMGB1 complex, similarly to HMGB1, induced neurite degeneration using an anti-pSer46-MARCKS antibody that detects MARCKS phosphorylation at Ser46, the original marker of neurite/synapse degeneration^30,45^ and with AT8 antibody detecting Tau phosphorylation at Ser202 and Thr205, a well-established marker of degeneration (Supplementary Fig.3). Western blotting revealed that dsHMGB1 at 0.56 nM, corresponding to the concentration of HMGB1 in CSF of MCI/AD patients^30^, but not atHMGB1 at the same concentration, increased pSer46-MARCKS (Supplementary Fig. 3a). As reported previously^45^, Aβ1-40 weakly affected MARCKS phosphorylation, although this did not reach statistical significance, while Aβ1-42 did not increase MARCKS phosphorylation (Supplementary Fig. 3a). dsHMGB1 but not atHMGB1, Aβ1-40 or Aβ1-42 increased Tau phosphorylation, as indicated by AT8 antibody (Supplementary Fig. 3b).

Contrastingly, when Aβ1-40 or Aβ1-42 formed the Aβ-dsHMGB1 complex with dsHMGB1, the resultant Aβ-dsHMGB1 complexes at 0.56 nM induced MARCKS phosphorylation at Ser46 (Supplementary Fig. 3a). However, when Aβ1-40 or Aβ1-42 complexed with atHMGB1, the Aβ-atHMGB1 complex at 0.56 nM did not induce MARCKS phosphorylation (Supplementary Fig. 3a). The neurodegenerative effects of Aβ1-40-dsHMGB1 and Aβ1-42-dsHMGB1 complexes were further confirmed by Western blot for AT8 that detects pathological Tau phosphorylation (Supplementary Fig. 3b).

We determined whether our original human monoclonal antibody against dsHMGB1 (human 129 antibody)^31^ affected the neurodegenerative effects of the Aβ1-40-dsHMGB1 or Aβ1-42-dsHMGB1 complexes (Supplementary Fig. 3c). The human 129 antibody at the same concentration (0.56 nM) suppressed both MARCKS and Tau phosphorylation induced by the dsHMGB1 and Aβ-dsHMGB1 complex (Supplementary Fig. 3c). siRNA knockdown of TLR4 similarly suppressed MARCKS and Tau phosphorylation induced by dsHMGB1 and the Aβ-dsHMGB1 complex (Supplementary Fig. 3d), suggesting that binding of the Aβ-dsHMGB1 complex to TLR4 mediated neurite degeneration. The results also suggested that the binding surfaces of Aβ1-40-dsHMGB1 and Aβ1-42-dsHMGB1 would be present on the outer domain(s) of dsHMGB1, and that Aβ1-40 and Aβ1-42 could stabilize the shapes of the binding domain(s).

Consistent with the effect of Aβ-dsHMGB1 complex on neurite degeneration, the Aβ-dsHMGB1 complex also induced cell death in mouse embryonic primary neurons at 0.56 nM, which is equivalent to the CSF HMGB1 concentration in MCI/AD patients^30^ (Supplementary Fig. 3e). The extent of cell death induction was approximately 70–80% of that induced by HMGB1 (Supplementary Fig. 3e). Though Aβ1-40 or Aβ1-42 alone did not induce cell death, both Aβ1-40 and Aβ1-42 acquired toxicity when they complexed with dsHMGB1, but not with atHMGB1 (Supplementary Fig. 3e).

Live imaging revealed that Aβ-dsHMGB1-induced cell death is morphologically characterized by ER ballooning, similar to the features of TRIAD present in early stages of AD in human patients and mouse models^30,31^ and consistent with our previous conclusion that secondary neuronal necrosis later forming Aβ plaques is also TRIAD^31^. Quantitative analyses confirmed that human 129 antibody at 0.56 nM also inhibited Aβ-dsHMGB1-induced cell death (Supplementary Fig. 3f).

Because surface plasmon resonance analysis was technically not suitable for evaluating binding of a protein complex to the receptor, we used an ELISA approach to evaluate the binding affinity of Aβ-dsHMGB1 complex to Toll-like receptor 4 (TLR4) (Supplementary Fig. 3g). The high affinities of Aβ1-40-dsHMGB1 (K_D_ = 3.816 x 10^-8^ M) and Aβ1-42-dsHMGB1 (K_D_ = 6.430 x 10^-8^ M) for TLR4, which were equivalent to that of dsHMGB1 (Supplementary Fig. 3g), supported the notion that binding of Aβ-dsHMGB1 complex to TLR4 induced neurite degeneration and cell death in the mechanisms reported previously, in which downstream TLR4 signaling induces neurite degeneration through MARCKS phosphorylation^45^ and cell death through Ku70 phosphorylation^30,31^.

To examine whether human monoclonal antibody against dsHMGB1 could inhibit binding of the Aβ-dsHMGB1 complex to TLR4, we pre-incubated the Aβ-dsHMGB1 complex and the anti-HMGB1 antibody (129) both at 1 nM for 30 min, and then analyzed the affinity of the Aβ-dsHMGB1 complex for TLR4 by ELISA (Supplementary Fig. 3h). As expected, the affinity of Aβ-dsHMGB1 complex to TLR4 after incubation with anti-dsHMGB1 antibody was suppressed significantly relative to Aβ-dsHMGB1 complex without pre-incubation (Supplementary Fig. 3h).

Collectively, these findings indicated that Aβ trapped with dsHMGB1 in dying neurons in the process of secondary necrosis results in formation of the Aβ-dsHMGB1 complex. In advanced stages of AD pathology, suppression of HMGB1 release by such intracellular formation of the Aβ-dsHMGB1 complex in neurons or by HMGB1 fixation to the Aβ-dsHMGB1 complex after incorporation of extracellular HMGB1 in microglia decreases extracellular HMGB1. On the other hand, extracellular Aβ-dsHMGB1 complex is increased by release from necrotic neurons. The Aβ-dsHMGB1 complex retained the toxicity like HMGB1 to induce synapse damage^30^ and secondary necrosis^31^. Although the affinity of the Aβ-dsHMGB1 complex for TLR4 is lower than that of dsHMGB1 for TLR4, the increased amount of extracellular Aβ-dsHMGB1 complex during advanced-stage of AD would promote AD pathology. Importantly, the toxicity of Aβ-HMGB1 complex was inhibited by our human anti-dsHMGB1 monoclonal antibody (CC129) (Supplementary Fig. 3e, f).

### Comparison of anti-HMGB1 and anti-Aβ antibodies on the toxicities of Aβ-HMGB1 complex

Given that some anti-Aβ antibodies including lecanemab are approved and used for human AD patients, and given that such antibodies might bind to the other surface of Aβ-dsHMGB1 complex, we next examined whether lecanemab could affect the interaction of Aβ-dsHMGB1 complex with TLR4 and whether it could influence the pathological changes of neurons by Aβ-dsHMGB1 complex.

First, we compared the inhibitory effect of lecanemab and that of our anti-dsHMGB1 antibodies on the TLR4 – Aβ-dsHMGB1 complex interaction by using ELISA, and revealed that two types of anti-dsHMGB1 antibody inhibited the TLR4 – Aβ-dsHMGB1 complex interaction more strongly than the anti-Aβ antibody lecanemab (Fig. 3a). Consistently, western blot analysis revealed that anti-dsHMGB1 antibodies at 0.56 nM inhibited MARCKS phosphorylation at Ser46 of normal iPSC-derived neurons cultured with 0.56 nM Aβ-dsHMGB1 complex in the medium far strongly than lecanemab at 0.56 nM (Fig. 3b). Suppression of pSer46-MARCKS signals by anti-dsHMGB1 antibodies but not by lecanemab in immunohistochemistry of normal iPSC-derived neurons cultured with 0.56 nM Aβ-dsHMGB1 complex also supported the conclusion from western blot analysis (Fig. 3c).

**Figure 3.**
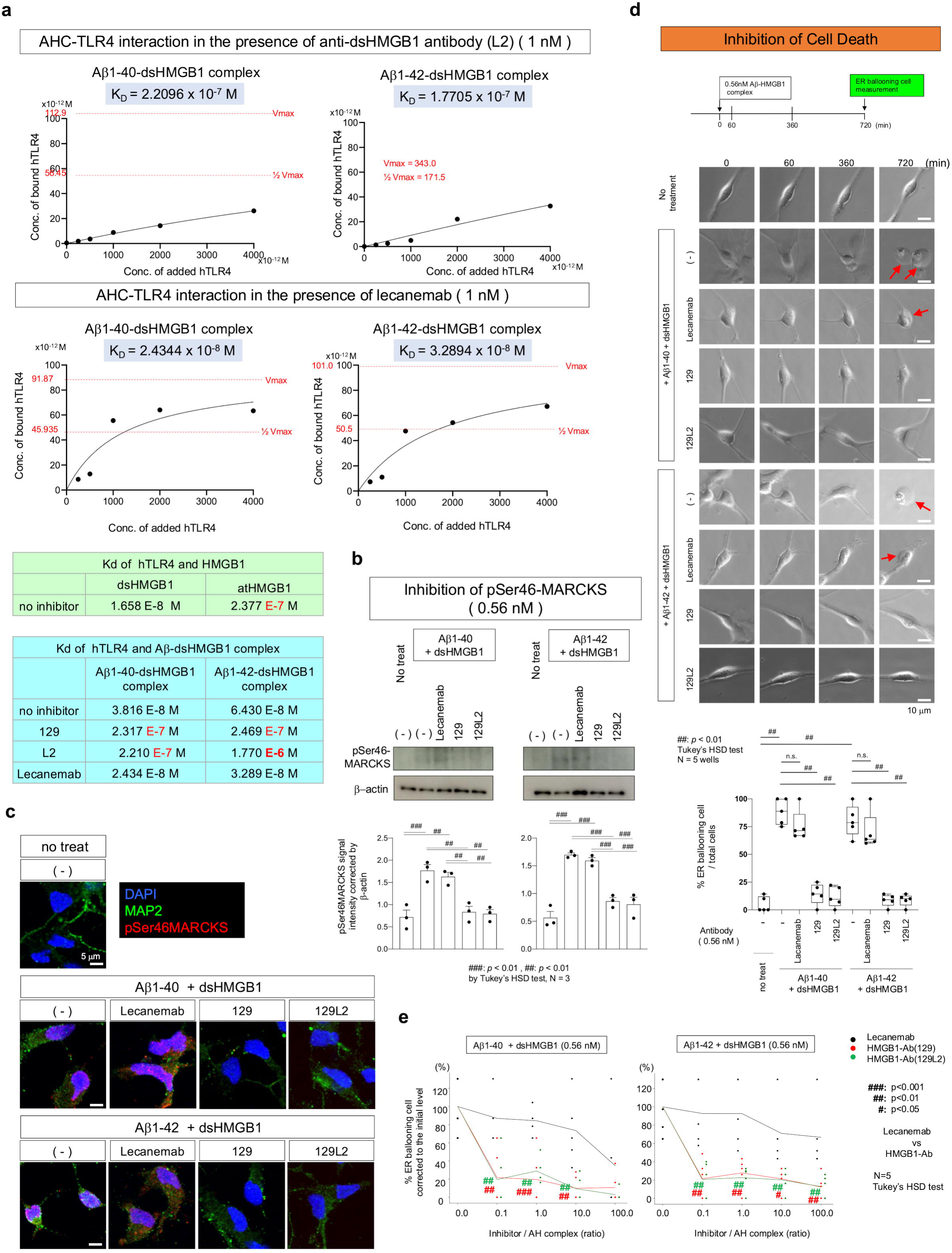
Suppressive effect of anti-HMGB1 antibody on Aβ-HMGB1 complex-induced neurodegeneration of human iPSC-derived neurons. a) Inhibition of the TLR4 – Aβ-dsHMGB1 complex interaction by two types of anti-dsHMGB1 antibody and anti-Aβ antibody was evaluated by ELISA. b) Western blot analysis of MARCKS phosphorylation at Ser46 of normal iPSC-derived neurons before and 30 min after addition of 0.56 nM Aβ-dsHMGB1 complex with or without 0.56 nM lecanemab, anti-dsHMGB1 antibody (CC129) or another anti-dsHMGB1 antibody (CC129L2). Lower graphs show intensities of the band signals corrected with GAPDH. **: p<0.01 in Tukey’s HSD test (N=3 wells per group). c) Immunohitochemistry of normal iPSC-derived neurons after addition of 0.56 nM of Aβ-dsHMGB1 complex with 0.56 nM lecanemab, anti-dsHMGB1 antibody (CC129) or another anti-dsHMGB1 antibody (CC129L2). **: p<0.01 in Tukey’s HSD test (N= 4 wells). d) Live imaging of normal human iPSC-derived neurons cultured during 720 min after addition of 0.56 nM Aβ-dsHMGB1 complex to the medium with one of the three antibodies. Upper panels show representative images of ER ballooning in neurons and lower graphs show the ratio of ER ballooning neurons to total neurons. Mean values of 5 visual fields from 3 wells were quantified. **: p<0.01 in Tukey’s HSD test (N=3 wells). e) The similar live imaging was performed with human iPSC-derived neurons in culture medium added with different concentrations of inhibitors (CC129, CC129L2 and lecanemab). Statistic differences are shown in the suppressive effect of neuronal death evaluated by ER ballooning. *: p<0.05, **: p<0.01, ***: p<0.001 in Tukey’s HSD test (N=3 wells).

Furthermore, we employed live imaging of normal iPSC-derived neurons cultured with 0.56 nM Aβ-dsHMGB1 complex in the medium to evaluate the Inhibition of Aβ-dsHMGB1 complex-induced neuronal death by three antibodies. Aβ-dsHMGB1 complex-induced neuronal cell death was remarkably suppressed by two types of anti-dsHMGB1 antibody (CC129 and CC129L2) but not by lecanemab when ER ballooning of neurons was evaluated at 720 min after addition of 0.56 nM Aβ-dsHMGB1 complex (Fig. 3d). By changing the concentration of inhibitors (CC129 antibody, CC129L2 antibody and lecanemab), live imaging further confirmed their differential suppressive effects on cell death of iPSC-derived neurons induced by Aβ-dsHMGB1 complex (Fig. 3e).

All these findings indicated predominance of anti-dsHMGB1 antibodies to an anti-Aβ-antibody lecanemab in the therapeutic effect on the toxicity of Aβ-dsHMGB1 complex.

### Diagnostic values of CSF- and plasma-Aβ-HMGB1 complex

These findings prompted us to examine the potential utility of Aβ-HMGB1 complex as a biomarker. To develop an Enzyme-Linked Immuno Sorbent Assay (ELISA) for measuring Aβ-HMGB1 complex as the total amount of its various molecular weight forms (Fig. 1), we selected the most appropriate anti-Aβ and anti-HMGB1 antibodies from commercially available ones to sandwich the Aβ-HMGB1 complex in an ELISA (Supplementary Fig. 4a), and used the ELISA assay to quantify Aβ-HMGB1 complex in CSF and plasma samples (Supplementary Fig. 4b, c) obtained from AD or MCI subjects and normal control subjects (Supplementary Table 1). Seventy CSF samples and 31 plasma samples from AD patients, 23 CSF and three plasma samples from MCI patients, and 48 CSF samples and 30 plasma samples from control subjects were enrolled in the study (Supplementary Table 1). The subject age, sex, MMSE, ADAS-cog, and CSF protein concentration are summarized in Supplementary Table 2.

Aβ-HMGB1 complex was markedly elevated in CSF and plasma of most AD patients (Supplementary Fig. 4b, c). Aβ-HMGB1 complex was not detected in most “clinically normal” control subjects, though in a few cases (NC11, 15, W232 in CSF; NC05, NC17 in plasma), who had not undergone amyloid-PET, exhibited high values in the CSF or plasma (Supplementary Fig. 4b, c). Among clinically normal control subjects with high Aβ-HMGB1 complex levels, only W232, revealing a below-normal Mini-Mental State Examination (MMSE) (24.0) score and elevated p-Tau (181 pg/mL), revisited the hospital and was assessed as early stage AD within 4 years.

In contrast to the elevated levels of HMGB1 in MCI patient CSF^30^, Aβ-HMGB1 complex was elevated in AD CSF but not MCI CSF (Supplementary Fig. 4b). The levels of HMGB1 and Aβ-HMGB1 complex in CSF collected from the same patients at the similar time revealed a negative relationship between HMGB1 and Aβ-HMGB1 complex in CSF (Supplementary Fig. 4d), suggesting that simultaneous measurement of CSF HMGB1 and Aβ-HMGB1 complex could be used for evaluating pathological stage of patients and for distinguishing between MCI and AD. Preliminarily, we examined the diagnostic capabilities of Aβ-HMGB1 complex with CSF and plasma samples from AD and MCI patients (Supplementary Fig. 4e, f). Receiver operating characteristic (ROC) analysis of comparisons between normal control subjects and AD or MCI subjects revealed that the area under the curve (AUC) was 0.7742 in AD vs. normal and 0.4667 in MCI vs. normal (Supplementary Fig. 4e), indicating that plasma levels of Aβ-HMGB1 complex could be used for the diagnosis of AD but not MCI. CSF levels of Aβ-HMGB1 were not able to distinguish AD or MCI patients from control subjects (Supplementary Fig. 4f), because the CSF Aβ-HMGB1 complex was not detectable in many patients.

We also addressed the relationship between cognitive function scores and plasma Aβ-HMGB1 complex levels (Supplementary Fig. 4g-j). Plasma Aβ-HMGB1 complex levels were positively correlated with ADAS-cog scores in a mixed population of AD patients (N = 30) and normal control subjects assessed clinically (N = 22) (Supplementary Fig. 4g). Moreover, annual change of ADAS-cog score in AD patients positively correlated with plasma Aβ-HMGB1 complex levels in patients that received sequential blood samplings, had detectable plasma Aβ-HMGB1 complex and received the ADAS-cog examination at multiple time points (N = 18) (Supplementary Fig. 4h). MMSE scores and plasma levels of Aβ-HMGB1 complex were also weakly negatively correlated in the same population of AD patients (N = 30) and normal control subjects diagnosed clinically (N = 22) (Supplementary Fig. 4i). Annual MMSE score change also correlated with plasma Aβ-HMGB1 complex levels in AD patients (N = 18) with detectable plasma Aβ-HMGB1 who had records of MMSE examination at multiple time points (Supplementary Fig. 4j).

### Combined analysis of Aβ-HMGB1 and HMGB1 stratified AD stages

Next, we examined whether Aβ-HMGB1 complex was able to stratify the pathological stages between MCI and AD. However, AUC values of CSF and plasma Aβ-HMGB1 complex were 0.600 and 0.725 respectively in ROC analyses, and insufficient to discriminate MCI from AD (Fig. 4a, b). Therefore, we assessed the predictive capability of combined use of CSF HMGB1 and Aβ-HMGB1 complex levels for improving the sensitivity and specificity (Fig. 4c, d, e). Though the increase was slight in combination of CSF Aβ-HMGB1 complex and CSF HMGB1 (0.644) (Fig. 4c), AUC values in stratification between MCI and AD became higher in combination of plasma Aβ-HMGB1 complex and CSF HMGB1 (0.758) or in combination of plasma Aβ-HMGB1 complex, CSF Aβ-HMGB1 complex and CSF HMGB1 (0.817) (Fig. 4d, e). These results supported that combined analysis of plasma Aβ-HMGB1 complex and CSF HMGB1 levels could potentially stratify MCI and AD (Fig. 4f), and suggested that it was superior to other markers using CSF-Aβ or Tau (Fig. 4f, g).

**Figure 4.**
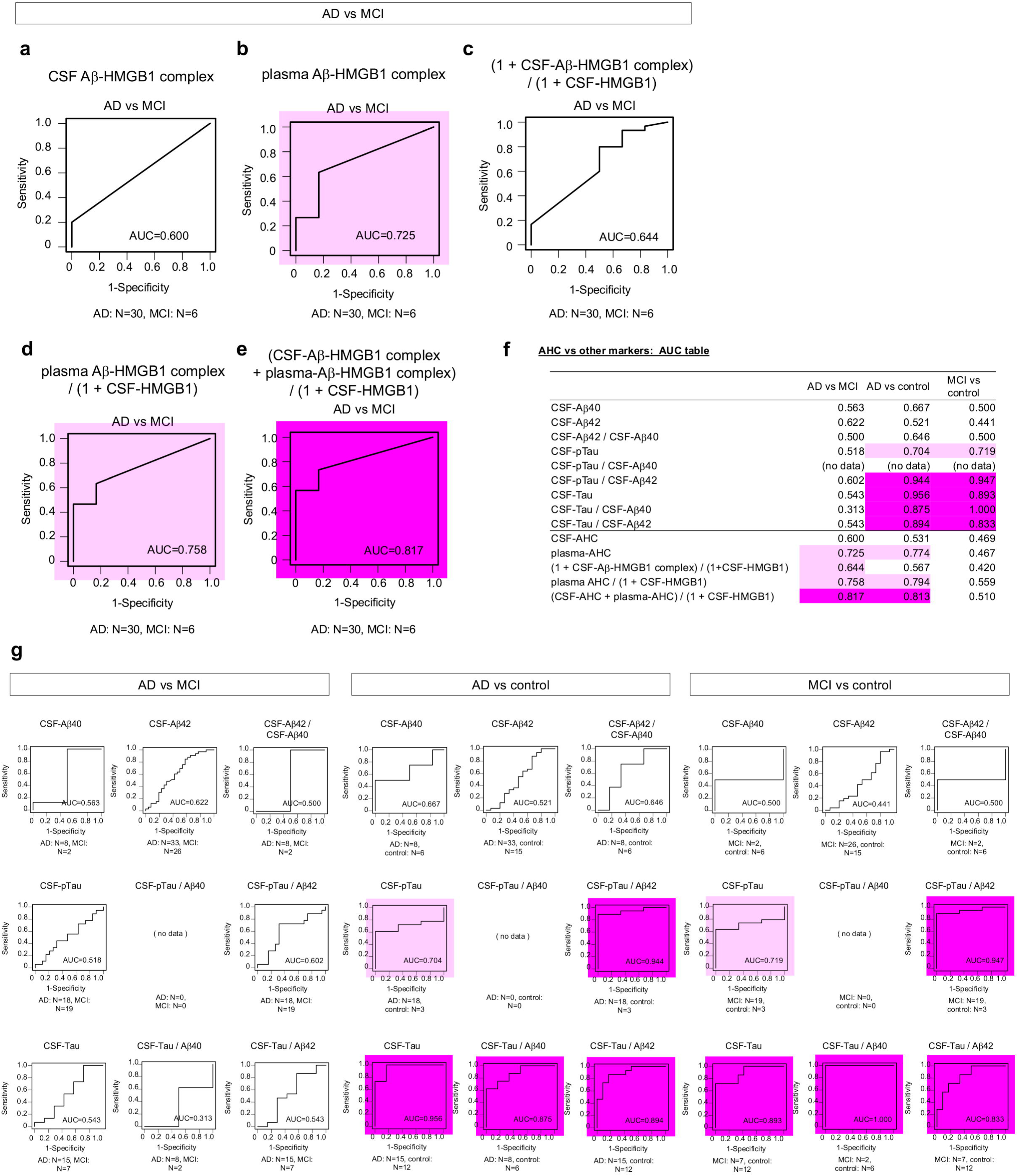
Analysis of Aβ-HMGB1 complex levels together with HMGB1 levels is useful for stratification of AD patients. a) ROC analysis to distinguish AD and MCI groups by CSF levels of Aβ-HMGB1 complex. b) ROC analysis to distinguish AD and MCI groups by plasma levels of Aβ-HMGB1 complex. c) ROC analysis to distinguish AD and MCI groups by CSF levels of Aβ-HMGB1 complex and CSF levels of HMGB1. d) ROC analysis to distinguish AD and MCI groups by plasma levels of Aβ-HMGB1 complex and CSF levels of HMGB1. e) ROC analysis to distinguish AD and MCI groups by using plasma Aβ-HMGB1 complex, CSF Aβ-HMGB1 complex and CSF HMGB1. f) Comparison of ROC analysis results by CSF-Aβ and/or CST-Tau or by Aβ-HMGB1 complex. Though currently available biomarkers are useful to distinguish between AD vs control or MCI vs control, they are not good at differential diagnosis between AD and MCI. Meanwhile, biomarkers based on Aβ-HMGB1 complex can distinguish between AD and MCI. Some of the AUC values are shown in Supplementary Fig 9. g) ROC analysis results to distinguish AD and MCI, AD and control or MCI and control groups by CSF-Aβ and/or CST-Tau that are currently used clinically. Light mazenda indicates AUC more than 0.7, and dark mazenda indicates AUC more than 0.8. Comparison a-f and g reveals a difference of Aβ-dsHMGB1 complex from previously established biomarkers such as pTau and Aβ that are useful for distinction between AD and control but not for distinction between AD and MCI.

### Therapeutic effects of anti-HMGB1 and anti-Aβ antibodies on Aβ-HMGB1 complex-related phenotypes

Finally, we compared the effects of anti-HMGB1 and anti-Aβ antibodies on the phenotypes of APP-KI (*App*^NL-G-F/NL-G-F^) mice^65^ carrying humanized *APP* mouse gene with Swedish (KM670/671NL), Beyreuther/Iberian (I716F) and Arctic (E693G) mutations. Y-maze and Morris water-maze (MWM) tests were performed to examine short-term and long-term spatial memories of post-onset APP-KI mice after a standard protocol treatment of lecanemab and treatments of a completely humanized 129 antibody (CC129L2)^66^ at various doses (Fig. 5a). In these experiments, APP-KI mice with saline injection and C57BL/6 mice were used as controls. The CC129L2 intravenous injections at 0.01, 0.02 and 0.2 mg/body weight kg every four weeks were effective both on cognitive functions of APP-KI mice in Y-maze and MWM tests, while lecanemab injected at 10 mg/ body weight kg every two weeks was not effective (Fig. 5a). We also tried to test 5xFAD mice to evaluate the effects of CC129L2 and lecanemab on cognitive functions, while 2 out of 3 lecanemab-treated mice died after 1^st^ or 2^nd^ injection due to the cerebral hemorrhage (Supplementary Fig. 5), and we could not confirm the effect on this model mice.

**Figure 5.**
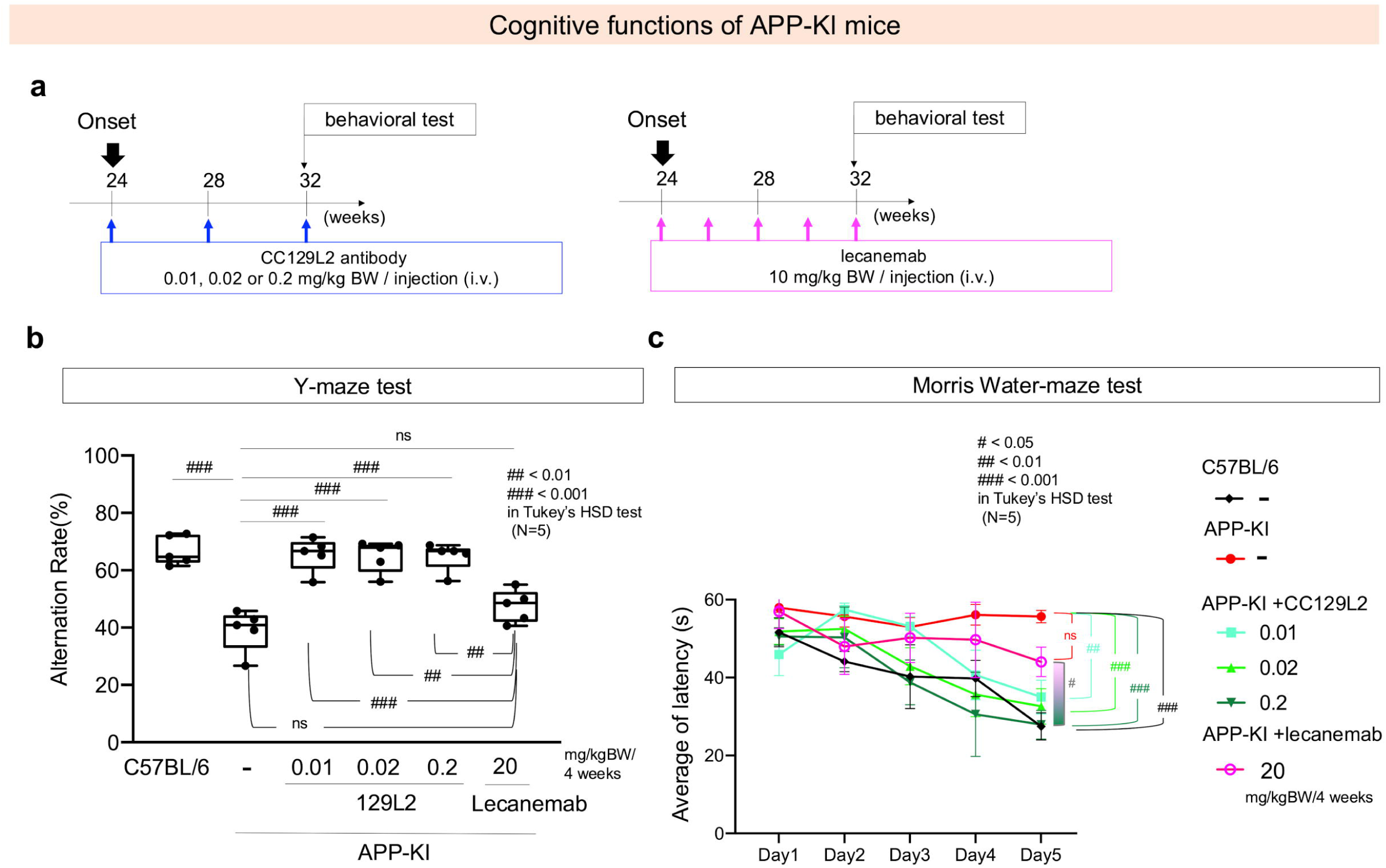
Comparison of therapeutic effects of anti-HMGB1 and anti-Aβantibodies on cognitive functions of AD model mice. a) Left and right panels show the protocols used for examining therapeutic effects of anti-HMGB1 (CC129L2) and anti-Aβ (lecanemab) antibodies, respectively. b) c) Left and right panels show results of APP-KI mice in Y-maze and Morris water maze tests respectively after a standard protocol treatment of lecanemab for human AD patients (10mg/BWkg biweekly) and treatments of CC129L2 at multiple doses (0.01, 0.02, 0.2 mg/BWkg every 4 weeks). #: p<0.05, ##: p<0.01, ###: p<0.001 in Tukey’s HSD test (N=5 mice).

Neuropathological examinations of the treated mice revealed reduction of extracellular Aβ aggregates both in the case of lecanemab and CC129L2 (Fig. 6). These findings were consistent with previous results of mAb158/lecanemab-treated mice that revealed reduction of Aβ aggregates but not improvement of cognitive functions^67,68^ and of CC129-treated mice that revealed both reduction of Aβ aggregates and improvement of cognitive functions^31^. Moreover, beyond previous results, we noticed critical findings in the pathology of APP-KI mice after treatments. The first difference between lecanemab-treated and CC129L2-treated mice was neuronal necrosis that was detected by positive staining of pSer46MARCKS and decreased signals of DAPI^30,31^. Neuronal necrosis was significantly reduced in CC129L2-treated APP-KI mice but not in lecanemab-treated APP-KI mice (Fig. 6). Unexpectedly, the second difference between lecanemab-treated and CC129L2-treated APP-KI mice was the amount of intraneuronal Aβ-HMGB1 complex (Fig. 6). Aβ-HMGB1 complex localized in neurons was significantly decreased in CC129L2-treated APP-KI mice but not in lecanemab-treated APP-KI mice (Fig. 6). Given that it was not plausible that the antibodies effectively remove substances within neurons, it might be the effect on neuronal vitality of C129L2-treatment removing extracellular HMGB1 and Aβ-HMGB1 complex both toxic to neurons that eventually led to increase intracellular amount of Aβ-HMGB1 complex. Our previous study indicated that intraneuronal accumulation of Aβ causes necrosis from early to late pathological stages^30,31^, and such accumulated intracellular Aβ at the late pathological stage could be actually that of Aβ-HMGB1 complex. The third difference was the ratio of nuclear and cytoplasmic HMGB1. HMGB1 signal intensities in the nuclei of neurons were recovered in CC129L2-treated mice but not in lecanemab-treated mice (Fig. 6) supported the recover of neuronal vitality by C129L2-but not by lecanemab-treatment. The fourth difference was the decrease of Aβ and HMGB-co-positive microglia clusters around small Aβ plaques, which were morphologically similar to pSer46MARCKS-positive neuronal necrosis by intracellular Aβ ^30^, in CC129L2-treated but not lecanemab-treated mice (Fig. 5b), suggesting that the reactive microglia to neuronal necrosis was suppressed by by C129L2-but not by lecanemab-treatment. Moreover, these effects on brain pathologies were also confirmed with CC129L2 intravenous injections at 0.01, 0.02 and 0.2 mg/body weight kg every four weeks (Supplementary Fig. 6). Collectively, these data suggested that CC129L2 could break such a negative chain of intracellular and extracellular Aβ-HMGB1 complex.

**Figure 6.**
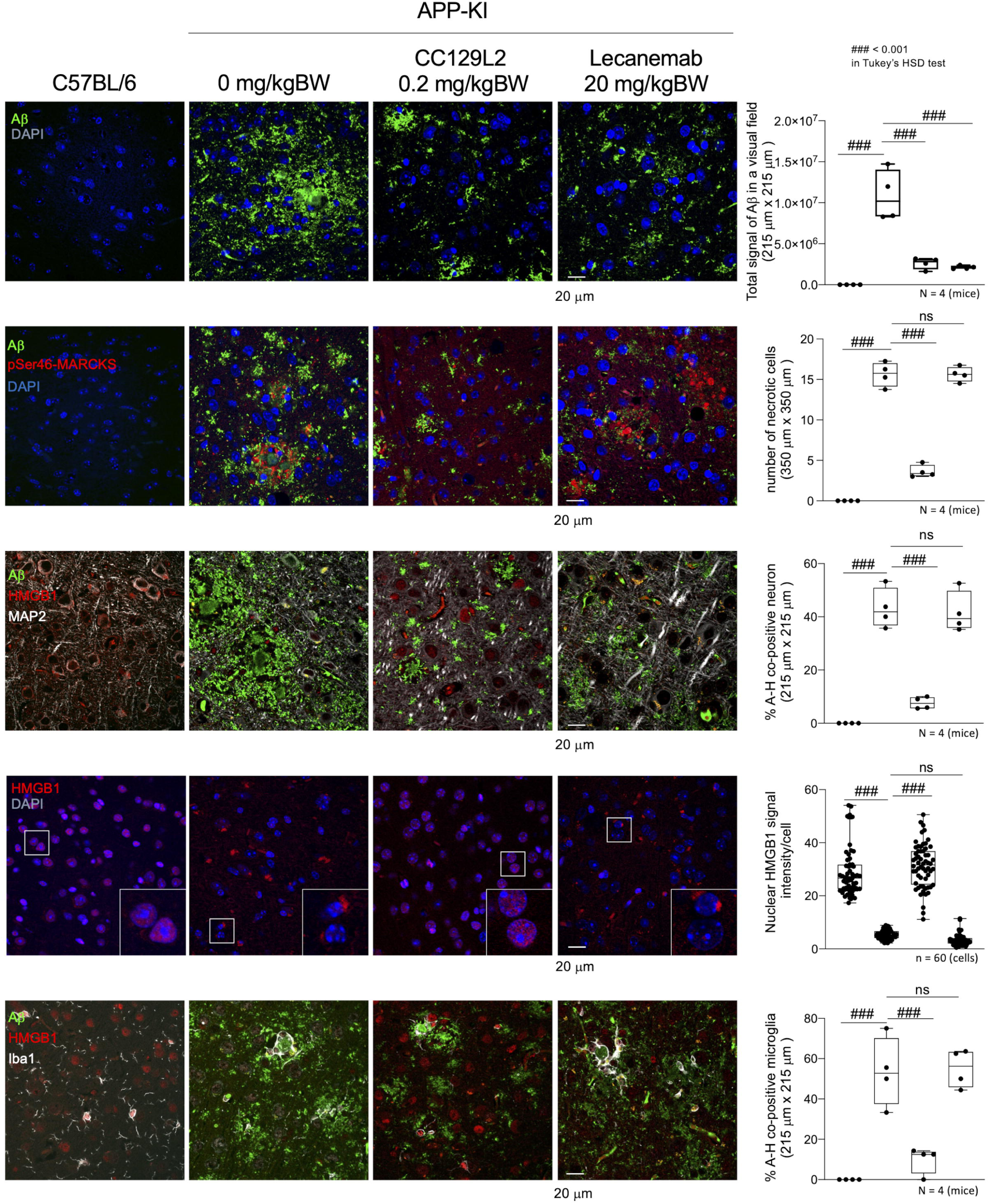
Therapeutic effects of anti-HMGB1 and anti-Aβ antibodies on neuropathology of APP-KI mice. Representative immunohistochemistry images for evaluation of Aβ burdens (Aβ, DAPI), necrotic neuronal death (Aβ, pSer46MARCKS, DAPI), Aβ-HMGB1 complex in neurons (Aβ, HMGB1, MAP2), cell viability evaluated by nuclear and cytoplasmic HMGB1 (HMGB1, DAPI) or Aβ-HMGB1 complex in microglia (Aβ, HMGB1, Iba1) are shown in C57BL/6 mice as the background mice of APP-KI mice, APP-KI mice after mock treatment of saline (0 mg/kgBW/4 weeks), CC129L2 (0.2 mg/kgBW/4 weeks) or lecanemab (20 mg/kgBW/4 weeks). Right graphs show quantitative comparisons of each factors with parietal cortex layer II-IV images obtained from 4 mice (mean value of 4 visual fields/mice) or with 60 cells (15 cells/mice x 4 mice) picked up randomly in the similar brain area for each group.

## Conclusion

This study reveals that Aβ-HMGB1 complex is a pathogenic molecule that promotes neurodegeneration mainly after the onset of AD, and suggests that it is also a biomarker detectable in the peripheral blood of AD patients whose elevation indicates an advanced stage of the patients. Immunoprecipitation analyses demonstrated the presence of Aβ-HMGB1 complex in dying neurons or activated neurons around Aβ plaques in human sporadic and familial AD brains, and immunohistochemical analyses revealed that Aβ-HMGB1 complex is generated in some organelles of neurons including Golgi apparatus and ER. Interaction of Aβ-HMGB1 complex with pattern recognition receptors such as TLR4 and its existence in microglia surrounding Aβ plaques suggest its relevance to brain inflammation. Interaction of Aβ-HMGB1 complex with TLR4 and its existence in dying neurons with ballooning ER and shrinking nuclei suggest that Aβ-HMGB1 complex induces TRIAD necrosis^30,57–59^, which is actually confirmed by human iPSC-derived neurons cultured with Aβ-HMGB1 complex. Its co-localization in neurons around Aβ plaques in 5xFAD and APP-KI mice at the advanced stages further supports that Aβ-HMGB1 complex is a pathogenic molecule to exponentially increase neuronal necrosis at advanced stages of AD pathology^30,31^. Collectively, the necrosis-enhancing function of Aβ-HMGB1 complex was similar to that of HMGB1, except that the pathological stages when they become active were different (Supplementary Fig. 7).

We also examine in this study the effect of our anti-dsHMGB1 antibodies and the approved anti-AD drug lecanemab by using human iPSC-derived neurons and AD model mice, and the results reveal that anti-dsHMGB1 antibodies but not lecanemab suppress the toxicity of Aβ-HMGB1 complex to induce neuronal necrosis. Recent results of clinical trials with lecanemab revealed statistically significant but still incomplete suppression (about -30%) of symptomatic progression (https://investors.biogen.com/news-releases/news-release-details/lecanemab-confirmatory-phase-3-clarity-ad-study-met-primary). The neuronal necrosis by Aβ-HMGB1 complex revealed in this study together with that by HMGB1 reported previously^30,31^ well explains the gap between symptomatic progression and suppressed Aβ-aggregate or Aβ-oligomer burdens of anti-Aβ−antibody-treated AD patients.

In the present study, we further reveal that Aβ-HMGB1 complex is detectable in peripheral blood. Among currently available biomarkers, neurofilament L (NFL) is of considerable significance, as circulating NFL levels are increased in multiple neurological diseases and correlate with the extent of neuronal cell death in multiple neurodegenerative diseases including AD^69^. However, NFL is not specific to AD and not linked to pathological stages of AD. Aβ-HMGB1 complex detectable in peripheral blood, especially combined with CSF-HMGB1, could complement such weak points of NFL and might be useful for clinical works.

## STAR METHODS

## KEY RESOURCE TABLE

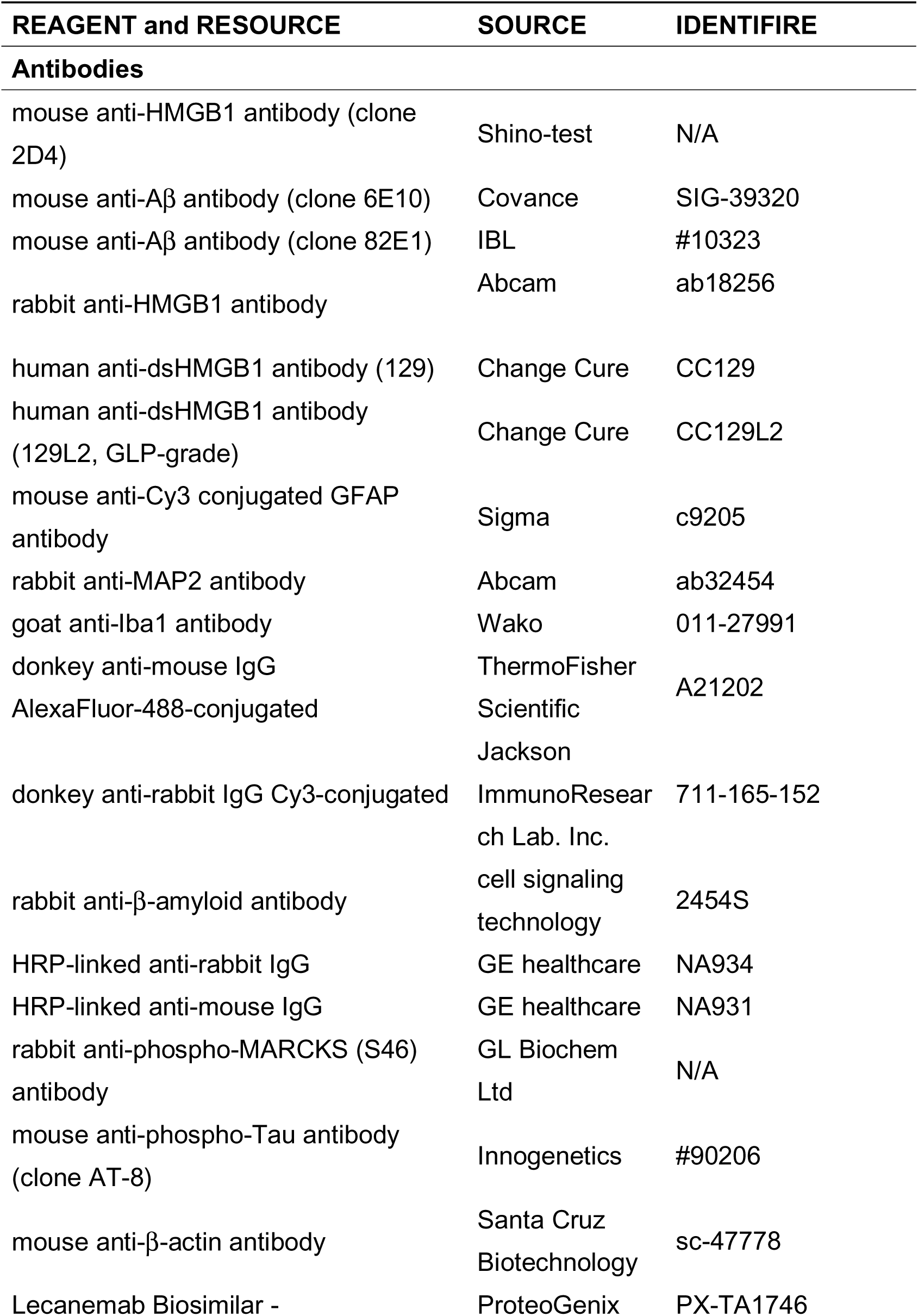
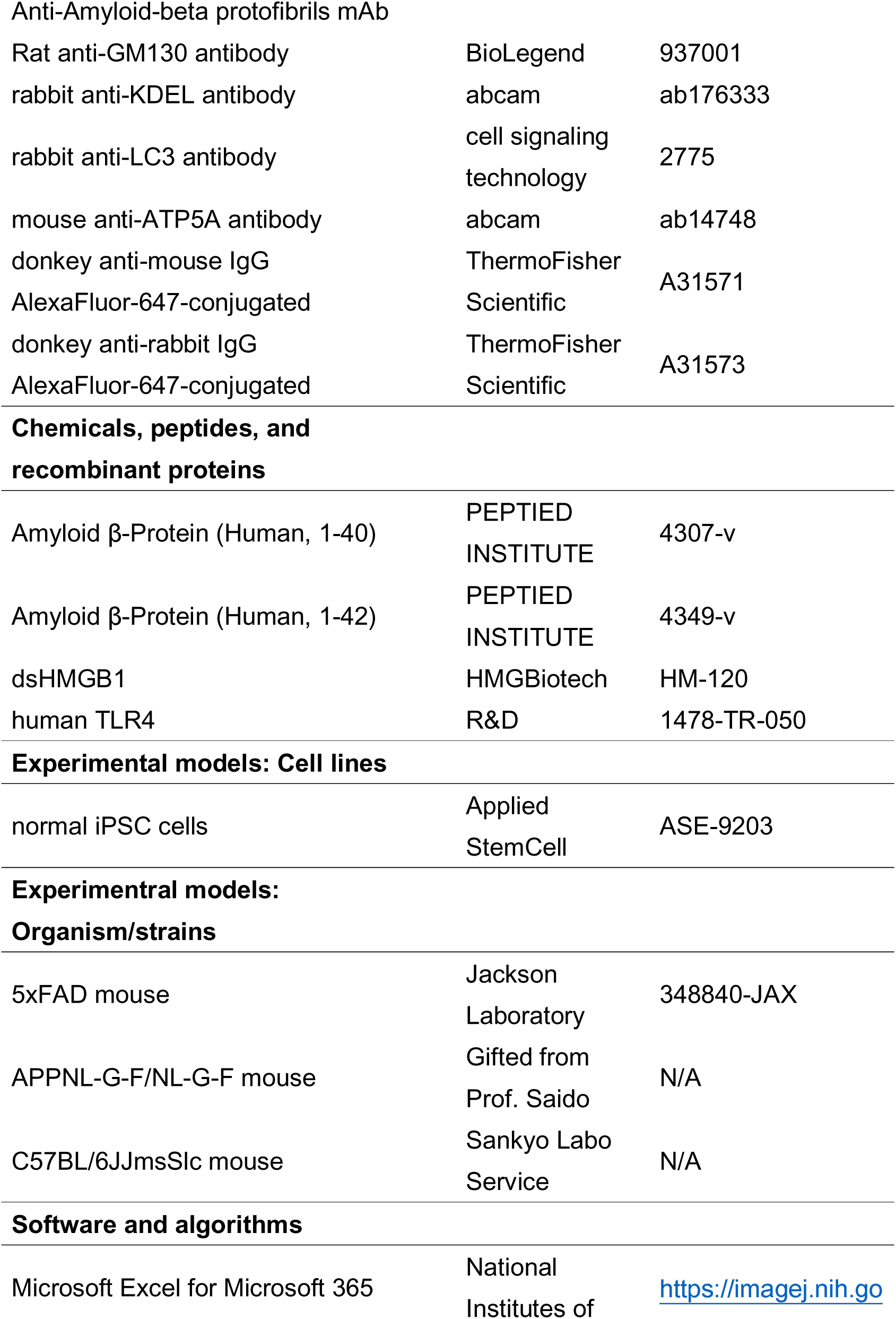
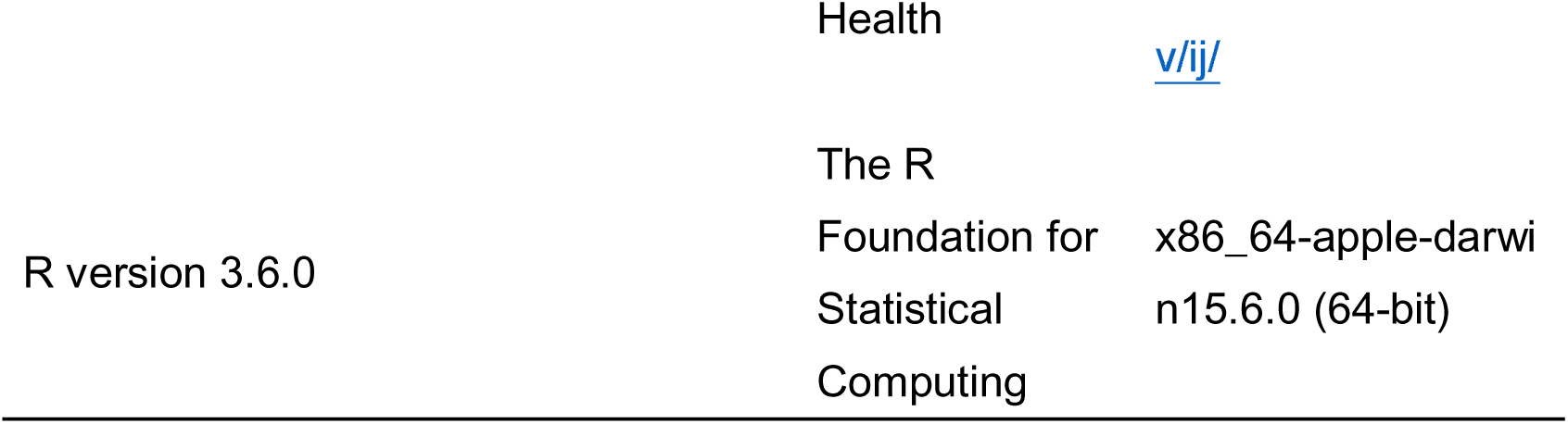

## Methods

### Aβ-HMGB1 complex ELISA

Ninety-six-well plates were coated with anti-HMGB1 monoclonal antibody (1 mg/L, clone 2D4, Shino-test, Tokyo, Japan) diluted in phosphate-buffered saline (PBS) and incubated overnight at 4°C. Unbound antibodies were removed by washing the plate with PBS containing 0.05% Tween 20 (washing buffer) and the remaining binding sites in the wells were blocked by incubating plates for 2 h with 400 μL/well PBS containing 1 % BSA at 37°C. After the plates were washed, human CSF or plasma samples or diluted calibrators (0, 175, 350, 700, 1400 pg/mL of Aβ-HMGB1 complex) were diluted to 1/4 concentrations with PBS containing 1% BSA, 100μL of each was to the well, and incubated for 20–24 hours at 37°C. Plates were washed and subsequently incubated with peroxidase-conjugated anti-Aβ antibody (mixture of clone 6E10 antibody (Previously Covance catalog# SIG-39320, Biolegend, San Diego, CA, USA) and clone 82E1 antibody (#10323, IBL, Gunma, Japan)) for 2 hours at room temperature. For detection, the substrate 3,3’,5,5’-tetra-methylbenzidine (Dojindo Laboratories) was added to each well. The reaction was terminated with 0.35 mol/L Na_2_SO_4_, and absorbance at 450 nm was measured on a microplate reader (model 680 Bio Rad, US). The working range for the assay was 30-2000 pg/mL.

### AD mouse models

Sagittal whole mouse brain sections were obtained from 5xFAD mice and APP-KI mice. 5xFAD transgenic mice overexpressing mutant human *APP* (770) with the Swedish (KM670/671NL), Florida (I716V), and London (V717I) familial Alzheimer’s disease (FAD) mutations and human *PS1* with FAD mutations (M146L and L285V) under the control of the mouse *Thy1* promoter were purchased from the Jackson Laboratory (34840-JAX, Bar Harbor, ME, USA). APP-KI (*App*^NL-G-F/NL-G-F^-KI) mice carrying a single human APP gene with the Swedish (KM670/671NL), Arctic (E693G), and Beyreuther/Iberian (I716F) mutations were a generous gift from Prof. Saido (Riken, Saitama, Japan).

### Immunohistochemistry

Mouse brains were fixed with 4% paraformaldehyde and embedded in paraffin. Sagittal sections (5 μm) were obtained using a microtome (REM-710, Yamato Kohki Industrial Co., Ltd., Saitama, Japan). Mouse and human sections were deparaffinized by xylene and ethanol incubation, and sections were boiled with 10 mM citrate buffer for 15 min for antigen retrieval. sections were boiled with 10 mM citrate buffer for 15 min for antigen retrieval. Sections were then incubated with 10% FBS diluted in PBS for 60 min and incubated with primary antibody for 12 hours at 4°C. The following primary antibodies were used: mouse anti-Aβ antibody (1:1000 or 1:100, #10323, IBL, Gunma, Japan), rabbit polyclonal anti-HMGB1 antibody (1:2000, ab18256, Abcam, Cambridge, UK), mouse anti-Cy3 conjugated GFAP (1:5000, c9205, sigma-Aldrich, St. Louis, MO, USA); rabbit anti-MAP2 (1:1000, ab32454, abcam, Cambridge, UK); and goat-anti-Iba1 (1:500, 011-27991, Wako, Osaka, Japan), mouse anti-ATP5A antibody (1:500, #ab14748, abcam, Cambridge, UK), rabbit anti-LAMP1 (1:50, #ab24170, abcam, Cambridge, UK), rabbit anti-KDEL (1:50, #ab176333, abcam, Cambridge, UK). For multiple co-staining, rabbit polyclonal anti-HMGB1 antibody (1:50, ab18256, abcam, Cambridge, UK) and mouse anti-Aβ antibody (1:100, #10323, IBL, Gunma, Japan) were labeled by Zenon Secondary Detection-Based Antibody Labeling Kits (Zenon Alexa Fluor 555 Rabbit IgG Labeling Kit, Z-25305, Thermo Fisher Scientific, Waltham, MA, USA) or (Zenon Alexa Fluor 488 Mouse IgG Labeling Kit, Z-25002, Thermo Fisher Scientific, Waltham, MA, USA). After washing sections with PBS, secondary antibodies were incubated for 1 hour at room temperature. The following secondary antibodies were used: donkey anti-mouse IgG AlexaFluor-488-conjugated (1:1000, A21202, Thermo Fisher Scientific, MA, USA), donkey anti-rabbit IgG Cy3-conjugated (1:1000, 711-165-152, Jackson Laboratory, CA, USA), donkey anti-mouse IgG AlexaFluor-647-conjugated (1:1000, A31571, Thermo Fisher Scientific, MA, USA), and donkey anti-rabbit IgG AlexaFluor-647-conjugated (1:1000, A31573, Thermo Fisher Scientific, MA, USA). After nuclear staining by incubation with 4′,6-diamidino-2-phenylindole (0.2 μg/mL in PBS, #D523, Dojindo Laboratories, Kumamoto, Japan), images were acquired using confocal microscopy (FV1200IX83, Olympus, Tokyo, Japan).

### Human CSF and plasma samples

Samples were collected from 71 AD patients, 26 MCI patients, and 48 normal subjects who had not been diagnosed with any brain diseases. Details are summarized in Supplementary Tables 1 and 2. Fasting CSF or plasma samples were obtained by lumbar puncture or blood sampling from elbow veins and collected into polypropylene tubes. CSF samples were centrifuged (1000 × *g* for 10 min at 4°C) to remove debris, and subsequently stored in small aliquots at -80°C. Blood samples were collected in tubes containing anti-coagulant (EDTA-2K) and centrifuged at 1,500*g* for 10 min at room temperature. Plasma samples were stored in small aliquots at -80°C.

#### Neuropsychological scores: Mini-Mental State Examination and ADAS-cog

The Japanese version of the Mini-Mental State Examination (MMSE) and Alzheimer’s Disease Assessment Scale-cognitive subscale 13 (ADAS-cog) were performed by the attending physician of each patient.

### Aβ and Tau measurement

CSF Aβ 1–40 and Aβ 1–42 were measured by ELISA using a human Aβ (1–40) ELISA kit (292-62301, Wako Chemical Co., Saitama, Japan) and human Aβ (1–42) ELISA kit (298-62401, Wako Chemical Co., Saitama, Japan). CSF p-Tau proteins were measured using INNOTEST Phospho-tau (181P, Innogenetics, Ghent, Belgium).

### Human tissue samples

Paraffin sections and frozen brain tissues were prepared from human MCI/AD brains and control brains without dementia (non-neurological disease controls). Informed consent for the use of human tissue samples was obtained after approval of the ethics committee at each institution and Tokyo Medical and Dental University.

### Preparation of A**β**-HMGB1 complex

Aβ peptide powder (Amyloid β-Protein (Human, 1-40), 4307-v, PEPTIDE INSTITUTE Osaka, Japan) (Amyloid β-Protein (Human, 1-42), 4349-v, PEPTIDE INSTITUTE, Osaka, Japan) was dissolved to 1 mM in DMSO. Disulfide HMGB1 (dsHMGB1, HMGBiotech, HM-120, Milano, Italy) was dissolved to 140 μM in PBS. Fully reduced HMGB1 (atHMGB1) was produced by incubating 100 μM dsHMGB1 with 20 mM DTT for 24 hours at 37°C. To compare atHMGB1 with dsHMGB1, 100 μM dsHMGB1 was also incubated in PBS for 24 hours at 37°C. These samples were then dialyzed at 4°C for 3 hours and replaced with PBS (Slide-A-Lyzer™ MINI Dialysis Device, Thermo Fisher Scientific, 69570, MA, USA).

HMGB1 and Aβ peptides were diluted to 60 μM in PBS, mixed at an equal molarity ratio, and incubated in a PCR thermal cycler with a heated lid for 48 hours at 37°C. The mixture was then dispensed in small quantities and immediately cooled to -80°C. When used in the experiments, Aβ-HMGB1 complexes were dissolved on ice and diluted with PBS to working concentrations prior to use.

### Tricin-SDS PAGE

Tricin-SDS PAGE (Schägger, 2006) was used to detect the Aβ-HMGB1 complex.

First, gels (Upper gel: 4%, Spacer gel: 10%, Lower gel: 16%) were cast 6 hours before electrophoresis. Next, an equal volume of sample buffer (1% SDS, 3% mercaptoethanol, 15% glycerol, 65 mM Tris/HCl (pH 7.0)) was added and the Aβ-HMGB1 complex was incubated at 37°C for 15 minutes and then electrophoresed in electrophoresis buffer (Anode buffer: 100 mM Tris, 22.5 mM HCl (pH 8.9), Cathode buffer: 100 mM Tris, 100 mM Tricin, 0.1% SDS (less than pH 8.25)) at 30 V for 30 min. The electrophoresis was then switched to 150V and stopped before the sample finished flowing. Gels were transferred onto Immobilon-P polyvinylidene difluoride membranes (Millipore, Burlington, MA, USA) using semi-dry transfer in electrode buffer (300 mM Tris, 100 mM acetic acid (pH 8.6)) at 0.4 mA /cm^2^ for 24 hours. Membranes were then incubated with primary and secondary antibodies after blocking with 5% skim milk in TBST (10 mM Tris-HCl, 150 mM NaCl, 0.05% Tween-20) for 1 hour. The following antibodies were used: rabbit anti-HMGB1 (1:2000, ab18256, Abcam, Cambridge, UK); rabbit-anti-β-amyloid antibody (1:1000, #2454S, Cell Signaling Technology, Danvers, MA, USA); mouse-anti-amyloid β (1:200, clone 82E1, #10323, IBL, Gunma, Japan); HRP-linked anti-rabbit IgG (1:3,000, NA934, GE Healthcare, Buckinghamshire, UK); and HRP-linked anti-mouse IgG (1:3,000, NA931, GE Healthcare, Buckinghamshire, UK). ECL Select Western Blotting Detection Reagent (RPN2235, GE Healthcare, Chicago, IL, USA) and a luminescence image analyzer were used to detect proteins (ImageQuant LAS 500, GE Healthcare, Chicago, IL, USA).

### Primary neuron culture

Mouse primary cortical neurons were obtained from E15 embryos. Four to Six total cerebral cortexes were dissected, incubated with 0.05% trypsin in 4 mL PBS (Thermo Fisher Scientific, #25200056, MA, USA) at 37°C for 15 min, then dissociated by pipetting. Cells were passed through a 70 μm cell strainer (Thermo Fisher Scientific, #22-363-548, MA, USA), collected by centrifugation, and cultured in neurobasal medium (Thermo Fisher Scientific, MA, USA) containing 2% B27, 0.5 mM L-glutamine, and 1% Penicillin/Streptomycin and 0.5 μM AraC. Seven days later, 0.56 nM (reflecting the CSF-HMGB1 concentration of a MCI/AD patient^30^ of dsHMGB1, atHMGB1, Aβ_1–40_, Aβ_1–42_, Aβ_1–40_-dsHMGB1 complex, Aβ_1–40_-atHMGB1 complex, Aβ_1–42_-dsHMGB1 complex, or Aβ_1–42_-atHMGB1 complex was applied to cells and incubated for 3 hours. For evaluation of the effect of human anti-HMGB1 antibody, stimulants (0.56 nM) were pre-incubated with the antibody (0.56 nM) for 30min at room temperature in medium, and the mixture was added to culture medium. Then, Western blotting was performed to assess neurite degeneration. To determine the effect of Aβ-HMGB1 complex on ER ballooning, time-lapse imaging of live cells was taken by BZ-X700 (Keyence, Osaka, Japan).

TLR4 knockdown was performed with TLR4 siRNA (Santa Cruz Biotechnology, Dallas, TX, USA) delivered with lipofectamine RNAiMAX reagent (#13778030, Thermo Fisher Scientific, MA, USA). Cells were incubated for 36 hours after siRNA knockdown prior to stimulant addition.

### Western Blot

Cultured cells were scraped and collected with PBS. After centrifugation (10000 *g* x 1 min at 4°C), cell pellets were lysed in extraction buffer containing 2% SDS, 1 mM DTT, and 10 mM Tris-HCl (pH 7.5). After heating at 100°C, lysates were centrifuged at 16000 *g* x 10 min at 4°C. Supernatants were then added to equal volumes of sample buffer containing 125 mM Tris-HCl (pH 6.8, Sigma, St. Louis, MO, USA), 4% SDS (Sigma, St. Louis, MO, USA), 20% glycerol (Wako, Osaka, Japan), 12% mercaptoethanol (Wako, Osaka, Japan), and 0.05% BPB (Nacalai, Kyoto, Japan). Samples were separated by SDS-PAGE, transferred onto Immobilon-P polyvinylidene difluoride membranes (Millipore, Burlington, MA, USA) using semi-dry transfer, and then blocked with 5% milk in TBST (10 mM Tris-HCl (pH 8.0, Sigma, St. Louis, MO, USA), 150 mM NaCl, 0.05% Tween-20). The antibodies listed below were suspended in Can Get Signal Solution (TOYOBO, Osaka, Japan) and incubated with membranes: rabbit anti-phospho-MARCKS (S46) (1:10,000, 1 hours at room temperature, GL Biochem Ltd, Shanghai, China); mouse anti-phospho-Tau (clone AT-8) (1:2,000, 3 hours at room temperature, #90206, Innogenetics, Gent, Belgium); mouse anti-β-actin (1:1000, 12 hour at 4 °C, #sc-47778, Santa Cruz Biotechnology, Dallas, TX, USA); mouse anti-TLR4 (1:5000, 3 hours at room temperature, #NB100-56566, Novus, CO, USA); HRP-linked anti-rabbit IgG (1:3,000, NA934, GE Healthcare, Buckinghamshire, UK); and HRP-linked anti-mouse IgG (1:3,000, NA931, GE Healthcare, Buckinghamshire, UK). ECL Select Western Blotting Detection Reagent (RPN2235, GE Healthcare, Chicago, IL, USA) and a luminescent image analyzer (ImageQuant LAS 500, GE Healthcare, Chicago, IL, USA) were used to detect proteins.

### ELISA for A**β**-HMGB1 complex and TLR4 affinity

For human anti-dsHMGB1 antibody treatment, 1 nM Aβ-HMGB1 complex and 1 nM antibody were incubated at room temperature for 30 min. For measurement, each type of Aβ-HMGB1 complex (1□nM in PBS, 100 μL) was added to 96-well plates pre-coated with HMGB1-Ab (HMGB1 ELISA kit Exp, 326078738, Shino-Test, Tokyo, Japan) and incubated for 24□h at 37°C. After plates were washed three times with wash buffer (HMGB1 ELISA KIT Exp, 326078738, Shino-Test, Tokyo, Japan), 100 μL peroxidase-labeled (Peroxidase Labeling Kit-NH2, LK11, Dojindo, Kumamoto, Japan) human TLR4 (1478-TR-050, R&D, Minneapolis, MN, USA) at 0, 0.25, 0.5, 1, 2 and 4 □nM in PBS was added to each well and incubated for 24 □hours at 25□°C. After washing the plates, 100□μL fluorescent reagent (HMGB1 ELISA KIT Exp, 326078738, Shino-Test, Tokyo, Japan) was added to each well and incubated for 30□min at 25°C. The reaction was stopped with 100□μL stop solution (HMGB1 ELISA KIT Exp, 326078738, Shino-Test, Tokyo, Japan) and absorbance was measured at 450 nm using a plate reader (SPARK 10M, TECAN, Grodig, Austria).

### iPSC differentiation to pan-neurons

Normal iPSCs (ASE-9203, Applied StemCell, CA, USA) were differentiated to pan-neurons as described previously^30^. Briefly, iPSCs were cultured in TeSR-E8 medium (STEMCELL Technologies, BC, Canada) with 10□μM Y27632 (253-00513, Wako, Osaka, Japan). After 24 h, medium was changed to Stem Fit (AK02N, Ajinomoto, Tokyo, Japan) containing 5□μM SB431542 (13031, Cayman Chemical, Ann Arbor, MI, USA), 5□μM CHIR99021 (13122, Cayman Chemical, Ann Arbor, MI, USA), and 5□μM dorsomorphin (044-33751, Wako, Osaka, Japan). After 5 days, iPS cells were dissociated with TrypLE Select (12563-011, Thermo Fisher Scientific, MA, USA). Neurospheres were then cultured in KBM medium (16050100, KHOJIN BIO, Saitama, Japan) with 20□ng/mL Human-FGF-basic (100-18B, Peprotech, London, UK), 10□ng/mL Recombinant Human LIF (NU0013-1, Nacalai, Kyoto, Japan), 10□μM Y27632 (253-00513, Wako, Osaka, Japan), 3□μM CHIR99021 (13122, Cayman Chemical, Ann Arbor, MI, USA), and 2□μM SB431542 (13031, Cayman Chemical, Ann Arbor, MI, USA) for 10 days. Finally, neurospheres were dissociated and seeded onto chambers coated with poly-L-ornithine (P3655, Sigma-Aldrich, St. Louis, MO, USA) and laminin (23016015, Thermo Fisher Scientific, Waltham, MA, USA). Pan-neurons were cultured in DMEM/F12 (D6421, Sigma-Aldrich, St. Louis, MO, USA) supplemented with 2% B27 (17504044, Thermo Fisher Scientific, Waltham, MA, USA), 1% Glutamax (35050061, Thermo Fisher Scientific, Waltham, MA, USA), and 1% penicillin/streptomycin (15140-122, Thermo Fisher Scientific, Waltham, MA, USA).

Five days later, 0.56 nM dsHMGB1, atHMGB1, Aβ_1–40_-dsHMGB1 complex, or Aβ_1–42_-dsHMGB1 complex was pre-incubated with 0.056 nM, 0.56 nM, 5.6nM, or 56nM Lecanemab Biosimilar - Anti-Amyloid-beta protofibrils mAb - (PX-TA1746, ProteoGenix, Schiltigheim, France), anti-dsHMGB1 antibody (CC129) or another anti-dsHMGB1 antibody (CC129L2) for 30min at room temperature in medium, and the mixture was added to culture medium containing NeuroFluor NeuO (ST-01801, STEMCELL Technologies, VC, Canada). After 720 minutes, the number of neurons with ER ballooning and the number of viable neurons were visually counted and the percentage of ER ballooning neurons to total neurons was determined from five images. Time-lapse imaging was taken by SS2000 (Yokogawa, Tokyo, Japan).

### Antibody therapeutics and cognitive function recovery

During the age from 6 to 8 months, APP-KI (AppNL-G-F/NL-G-F) mice or C57BL/6J mice received injection into the tail vein of the human monoclonal anti-HMGB1 antibody (CC129L2, Change Cure, Kanagawa, Japan) at 0.2□, 0.02 or 0.01 mg/kgBW once a month, or of the human monoclonal anti-Amyloid β antibody (Lecanemab Biosimilar, ProteoGenix, Schiltigheim, France) at 10 mg/kgBW once every two weeks. Y-maze test and Morris water maze test were performed at 8 months of age. In Y-maze test, Y-shape maze consisting of three identical arms with equal angles between each arm (O’HARA & Co., Ltd, Tokyo, Japan) was used. Mice was placed at the end of one arm and allowed to move freely through the maze during an 8□min session. The percentage of spontaneous alterations (indicated as an alteration score) was calculated by dividing the number of entries into a new arm that was different from the previous one with the total number of transfers from an arm to another arm. In Morris water maze test, mice performed four trials (60 sec) per day for 5 days, and the latencies to reach the platform were measured. After behavioral tests, the mouse brain, heart, lung, liver, spleen, kidney, intestine, muscle, skin and spinal cord were dissected, and the organ tissues were fixed with 4% paraformaldehyde for 24 hours, washed with PBS, then embedded in paraffin. The paraffin sections were used for immunohistochemistry or hematoxylin-eosin staining.

### Statistics

Statistical analyses for human biological examinations were performed using R version 3.6.2. A box plot was used to depict data distribution, and the data are also plotted as dots. Box plots show medians, quartiles, and whiskers, which represent data outside the 25^th^–75^th^ percentile range. p-values were calculated using Wilcoxon’s rank sum test with post-hoc Bonferroni correction.

Student’s t-test was used for comparisons between groups in the other biological experiences. For multiple group comparison, Tukey’s HSD test was employed.

Correlation analysis was performed using Microsoft Excel for Microsoft 365, and the p-value for Pearson’s correlation coefficient was calculated using R. Receiver operating characteristic (ROC) was analyzed using the ROCR package for R.

## Study approval

This study was performed in strict accordance with the Guidelines for Proper Conduct of Animal Experiments by the Science Council of Japan. This study was approved by the Committees on Gene Recombination Experiments, Human Ethics, and Animal Experiments of the Tokyo Medical and Dental University (G2018-082C3, O2020-002-03, and A2021-211A, respectively).

## Author Contributions

H.H., Y.Y., K.F., S.Y.: data acquisition and analysis and drafting of the manuscript. K.F., A.I., N.A., M.W., T.M., M.K., G.S., H.A., A.I.: patients sample acquisition. H.T. and H.O.: conception and design of the study, obtaining funding, and drafting the manuscript.

## Acknowledgment

This work was supported by grants to H.O., including a Grant-in-Aid for Scientific Research on Innovative Areas (Foundation of Synapse and Neurocircuit Pathology, 22110001/ 22110002) from the Ministry of Education, Culture, Sports, Science, and Technology of Japan (MEXT), and a Grant-in-Aid for Scientific Research A (16H02655, 19H01042, 22H00464) from the Japanese Society for the Promotion of Science (JSPS).

## Data Availability

All data generated or analyzed during this study are included in this article. Source data are provided with the paper.

## Consent to participate

Informed consent was obtained from all individual participants included in the study.

## Figure Legends

**Supplementary Figure 1**

**Aβ-HMGB1 complex is generated in necrotic cells surrounding Aβ plaques in AD model mice and human AD patients**

a) Immunohistochemistry with anti-Aβ and HMGB1 antibodies of 5xFAD mouse cortex (parietal cortex) at 6 months old. Yellow arrowheads indicate co-staining with HMGB1 (red) and Aβ (green) in the cytoplasm of neurons surrounding Aβ plaques. In these neurons, HMGB1 was shifted from the nucleus to the cytoplasm. In addition, a ballooning cell was observed in, which HMGB1 (red) and Aβ (green) signals were pushed out to the periphery of the cytoplasm just before rupture (yellow arrow). In normal neurons, HMGB1 remained in the nucleus and Aβ was not accumulated in the cytoplasm (white arrow). The lower graph shows quantitative analysis and statistical analysis (Student’s t-test) of cells with Aβ and HMGB1 co-localization. See also Supplementary Video 1.

b) Immunohistochemistry with anti-Aβ and HMGB1 antibodies of APP-KI mouse cortex (parietal cortex) at 6 months old. Similar to 5xFAD mice, co-staining of HMGB1 (red) and Aβ (green) was present in the cytoplasm of abnormal neurons surrounding Aβ plaques (yellow arrowhead), while in normal neurons, HMGB1 remained in the nucleus and Aβ was not accumulated in the cytoplasm (white arrow). The lower graph shows quantitative analysis and statistical analysis (Student’s t-test) of cells with Aβ and HMGB1 co-localization. See also Supplementary Video 2.

c) Immunohistochemistry for Aβ and HMGB1 of human postmortem cerebral cortex sections of familial AD patients (PS1-M146L). Two different images are shown. In both images, representative neurons are denoted with squares. HMGB1 was shifted from the nucleus to the cytoplasm, and Aβ and HMGB1 co-localized in the cytoplasm (#1 and #2). In advanced AD stages (#3, #4), nuclear DAPI signal was weakened, halo-like ballooning of the cytoplasm (v) occurred, and shrinkage of residual cell bodies followed. These changes are typical morphological features of TRIAD necrosis. In final stage AD (#4), cytoplasmic ballooning ruptured and Aβ-HMGB1 complex was released from cells, consistent with live imaging of TRIAD necrosis in primary neurons^30,58^. The white arrow indicates a normal neuron with high nuclear HMGB1 signal, and the blue dot arrow indicates a capillary blood vessel. Right graph shows quantitative analysis and statistical analysis (Student’s t-test) of cells with Aβ and HMGB1 co-localization. See also Supplementary Video 3, 4.

d) Immunohistochemistry for Aβ and HMGB1 of human postmortem cerebral cortex sections of sporadic AD patients. #1 and 2 are representative neurons whose cytoplasm revealed co-localized Aβ and HMGB1.

e) Image of co-localization of Aβ and HMGB1 signals surrounding a large Aβ plaque in the cerebral cortex of 5xFAD mice at 6 months of age. See also Supplementary Video 5.

f) Co-localization of Aβ and HMGB1 signals surrounding a large Aβ plaque in the cerebral cortex of APP-KI mice at 6 months of age. See also Supplementary Video 6.

g) Immunostaining of neuron-specific marker MAP2 in the parietal cortex of 5xFAD mice at 6 months of age. Yellow arrows indicate neurons with cytoplasmic Aβ and HMGB1 co-localization.

h) Immunostaining of neuron-specific marker MAP2 in the parietal cortex of APP-KI mice at 6 months of age. Yellow arrows indicate neurons with cytoplasmic Aβ and HMGB1 co-localization.

i) Immunostaining of neuron-specific marker MAP2 in the parietal lobe cortex of a human PS1-linked AD patient. The left panels show low-magnification images. Neurons with Aβ and HMGB1 co-localization are indicated (yellow arrows). Right panels show high-magnification images of two regions (#1, #2) including neurons with Aβ and HMGB1 co-localization (yellow arrows). These neurons exhibited cytoplasmic leakage of HMGB1 (#1, #2), faint DAPI staining (#2), and cytoplasmic vacuoles (#1, #2) characteristic of TRIAD necrosis. The neurons did not exhibit features of apoptosis such as chromatin condensation or apoptotic bodies.

j) Immunostaining of neuron-specific marker MAP2 in the parietal lobe cortex of a sporadic AD patient. Lower panels show high-magnification images of two representative regions (#1, #2) including neurons with Aβ and HMGB1 co-localization (yellow arrows).

**Supplementary Figure 2**

**Aβ-HMGB1 complex is scavenged by microglia but not detected in astrocytes**

Immunohistochemistry of cell type-specific markers was performed for identification of the Aβ-HMGB1 complex generation site *in vivo*. Results from immunostaining with the neuron-specific marker MAP2 are shown in Fig. 4. Here immunostainings with the microglia-specific marker Iba1 and astrocyte-specific marker GFAP are shown.

a) Immunostaining with the microglia-specific marker Iba1 in the parietal cortex of 5xFAD mice at 6 months of age. Yellow arrows indicate merged Aβ and HMGB1 signals in microglia.

b) Immunostaining of the microglia-specific marker Iba1 in the parietal cortex of APP-KI mice at 6 months of age. Yellow arrows indicate merged Aβ and HMGB1 signals in microglia.

c) Immunostaining with the microglia-specific marker Iba1 in the parietal lobe cortex of a human PS1-linked AD patient. Left panels show low-magnification images. Right panels show high-magnification images of two regions (#1, #2) including microglia. Yellow arrows indicate merged Aβ and HMGB1 signals in microglia.

d) Immunostaining with the astrocyte-specific marker GFAP in the parietal cortex of 5xFAD mice at 6 months of age. Light blue arrows indicate astrocytes that do not contain merged Aβ and HMGB1 signals.

e) Immunostaining with the astrocyte-specific marker GFAP in the parietal cortex of APP-KI mice at 6 months of age. Light blue arrows indicate astrocytes that do not contain merged Aβ and HMGB1 signals.

f) Immunostaining with the astrocyte-specific marker GFAP in the parietal lobe cortex of a human PS1-linked AD patient. Left panels show low-magnification images. Right panels show high-magnification images of two regions (#1, #2) including astrocytes or astrocyte processes (#3). Though the #1 astrocyte did not contain merged Aβ and HMGB1 signals, the #2 astrocyte with an enlarged cytoplasm, which is reminiscent of reactive astrocytes/gemistocytes, contained merged Aβ and HMGB1 signals (yellow arrow). Astrocyte process #3 also contained merged Aβ and HMGB1 signals (yellow arrow).

**Supplementary Figure 3**

**Aβ-HMGB1 complex induces neurite degeneration and neuronal death Via binding to Toll-like receptor 4**

a) Western blotting revealed that Aβ-HMGB1 complex induced phosphorylation of MARCKS at Ser46 (pSer46-MARCKS). Primary mouse cortical neurons (E15) were incubated for 3 hours with monomeric disulfide HMGB1 (dsHMGB1), monomeric full-reduced-/all-thiol-HMGB1 (atHMGB1), Aβ 1-40/1-42 monomer immediately after dissolution of the powder, Aβ-dsHMGB1 complex, or Aβ-atHMGB1 complex (0.56 nM). Aβ-dsHMGB1 complex and dsHMGB1 increased pSer46-MARCKS. The bottom graph shows quantitative analysis and statistical analysis (Tukey’s HSD test). N=5 wells per group.

b) Western blotting with anti-Tau antibody (AT-8) revealed the Tau-phosphorylating effects of dsHMGB1 monomer, atHMGB1 monomer, Aβ 1-40/1-42 monomer, and their complexes with dsHMGB1 or atHMGB1. Monomeric dsHMGB1, Aβ1-40-dsHMGB1 complex, and Aβ1-42-dsHMGB1 complex induced Tau phosphorylation. The bottom graph shows quantitative analysis and statistical analysis (Tukey’s HSD test). N=5 wells per group.

c) A monoclonal antibody against human dsHMGB1 (0.56 nM) inhibited dsHMGB1-induced and Aβ-HMGB1 complex-induced phosphorylation of MARCKS and Tau. The bottom graphs show quantitative analysis and statistical analysis (Tukey’s HSD test). N=5 wells per group.

d) siRNA knockdown of TLR4 similarly inhibited MARCKS and Tau phosphorylation. The bottom graphs show quantitative analysis and statistical analysis (Tukey’s HSD test). N=5 wells per group.

e, f) The upper panel shows the protocol for cell death analysis of primary cortical neurons (E15) treated with Aβ-HMGB1 complex. The middle panels show dsHMGB1-induced or Aβ-dsHMGB1 complex-induced cell death (white arrows) with ER ballooning (right panels), a typical feature of TRIAD necrosis, in cortical neurons observed by live imaging of primary cultures. The lower graph shows quantitative analysis of TRIAD necrosis induced by Aβ-HMGB1 complex before

(e) or after (f) pre-incubation with human anti-human HMGB1 antibody (0.56 nM). Human anti-human HMGB1 antibody (#129) inhibited TRIAD necrosis induced by Aβ-HMGB1 complex. N (number of wells) = 5. #: p<0.05, ##: p<0.01, Tukey’s HSD test.

g) ELISA for binding of Aβ-HMGB1 complex to TLR4. Disulfide HMGB1 (dsHMGB1)-Aβ complex, but not full-reduced HMGB1 (redHMGB1)-Aβ complex, exhibited dose-dependent TLR4 binding.

h) Human anti-human HMGB1 antibody blocked binding of Aβ-HMGB1 complex to TLR4.

**Supplementary Figure 4**

**Diagnostic ability of CSF and plasma Aβ-HMGB1 complex for AD**

a) Schematic of ELISA method for detection of Aβ-HMGB1 complex.

b) Concentrations of CSF Aβ-HMGB1 complex in the control group (N □= □48 subjects), MCI group (N□ =□ 23 subjects), and AD group (N□ =□ 70 subjects) as measured by ELISA. Although CSF Aβ-HMGB1 complex was elevated in the AD group relative to the MCI and control groups, the value was exceptionally high in three cases of the control group (NC11, NC15, W232). **: p<0.01, Wilcoxon’s rank sum test with post-hoc Bonferroni correction.

c) Plasma Aβ-HMGB1 complex concentrations in the control group (N = 30 subjects), MCI group (N = 3 subjects), and AD group (N = 31 subjects) were measured by ELISA. Two cases of the control group had high values. CSF and plasma of these exceptional cases in Fig. 1b and c were not sampled on the same date. **: p<0.01, Wilcoxon’s rank sum test with post-hoc Bonferroni correction.

d) Parallel coordinate plots showing the relationship between CSF HMGB1 and Aβ-HMGB1 complex.

e) ROC analysis of sensitivity and specificity to distinguish AD and control, MCI and control, or AD and MCI groups by plasma levels of Aβ-HMGB1 complex.

f) ROC analysis of sensitivity and specificity to distinguish AD and control, MCI and control, or AD and MCI groups by CSF levels of Aβ-HMGB1 complex.

g) Correlation between plasma Aβ-HMGB1 complex levels and ADAS-cog

scores.

h) Correlation between plasma Aβ-HMGB1 complex levels and annual changes of ADAS-cog scores.

i) Correlation between plasma Aβ-HMGB1 complex levels and MMSE scores.

j) Correlation between plasma Aβ-HMGB1 complex levels and annual changes of MMSE scores. *r* denotes Pearson’s correlation coefficients.

**Supplementary Figure 5**

**Pathological findings of 5xFAD mice died after treatment of lecanemab**

Two among three mice died after 1^st^ or 2^nd^ administration of lecanemab (10 mg/kgBW/2 weeks). Upper panel shows hematoxylin-eosin staining of cerebral cortex of a 5xFAD mouse with no treatment. Lower panel shows the similar staining of a 5xFAD mouse died two days after 1^st^ administration. Blood congestion and micro-hemorrhage foci were observed. Upper and lower right panels show macroscopic findings of mouse brains. Whole brain surface of the 5xFAD mouse that died after lecanemab administration was red.

**Supplementary Figure 6**

**Therapeutic effects of different doses of anti-HMGB1 antibody on neuropathology of APP-KI mice**

Representative immunohistochemistry images for evaluation of Aβ burdens (Aβ, DAPI), necrotic neuronal death (Aβ, pSer46MARCKS, DAPI), Aβ-HMGB1 complex in neurons (Aβ, HMGB1, MAP2), cell viability evaluated by nuclear and cytoplasmic HMGB1 (HMGB1, DAPI) or Aβ-HMGB1 complex in microglia (Aβ, HMGB1, Iba1) are shown in APP-KI mice after mock treatment of saline (0 mg/kgBW/4 weeks) and CC129L2 (0.01, 0.02 or 0.2 mg/kgBW/4 weeks). Right graphs show quantitative comparisons of each factors. Right graphs show quantitative comparisons of each factors with parietal cortex layer II-IV images obtained from 4 mice (mean value of 4 visual fields/mice) or with 60 cells (15 cells/mice x 4 mice) picked up randomly in the similar brain area for each group.

**Supplementary Figure 7**

**Two biomarkers and one targeted therapy covers early and late AD stages**

a) Relationship between clinical AD stages and two biomarkers.

b) Binding of HMGB1 and Aβ-dsHMGB1 complex to TLR4, and their suppression by anti-HMGB1 antibody.

**Supplementary Table 1: Summary of study cohort clinical information**

**Supplementary Table 2: Clinical information of individual patients in the study cohort**

**Supplementary Video 1: 3D images of double staining of Aβ and HMGB1 in neurons surrounding Aβ plaques in 5xFAD mice**

**Supplementary Video 2: 3D images of double staining of Aβ and HMGB1 in neurons surrounding Aβ plaques in *App*^NL-G-F/NL-G-F^-KI mice**

**Supplementary Video 3: 3D images of double staining of Aβ and HMGB1 in a human AD brain.**

**Supplementary Video 4: 3D images of double staining of Aβ and HMGB1 in a human AD brain**

**Supplementary Video 5: 3D images of double staining of Aβ and HMGB1 in neurons surrounding Aβ plaques in 5xFAD mice**

**Supplementary Video 6: 3D images** of double staining of Aβ and HMGB1 in neurons surrounding Aβ plaques in *App*^NL-G-F/NL-G-F^-KI mice

