## Supplementary figures and images for "Aβ-HMGB1 complex is a pathogenic molecule at the advanced stage of Alzheimer’s disease"

### Supplementary Figure 1

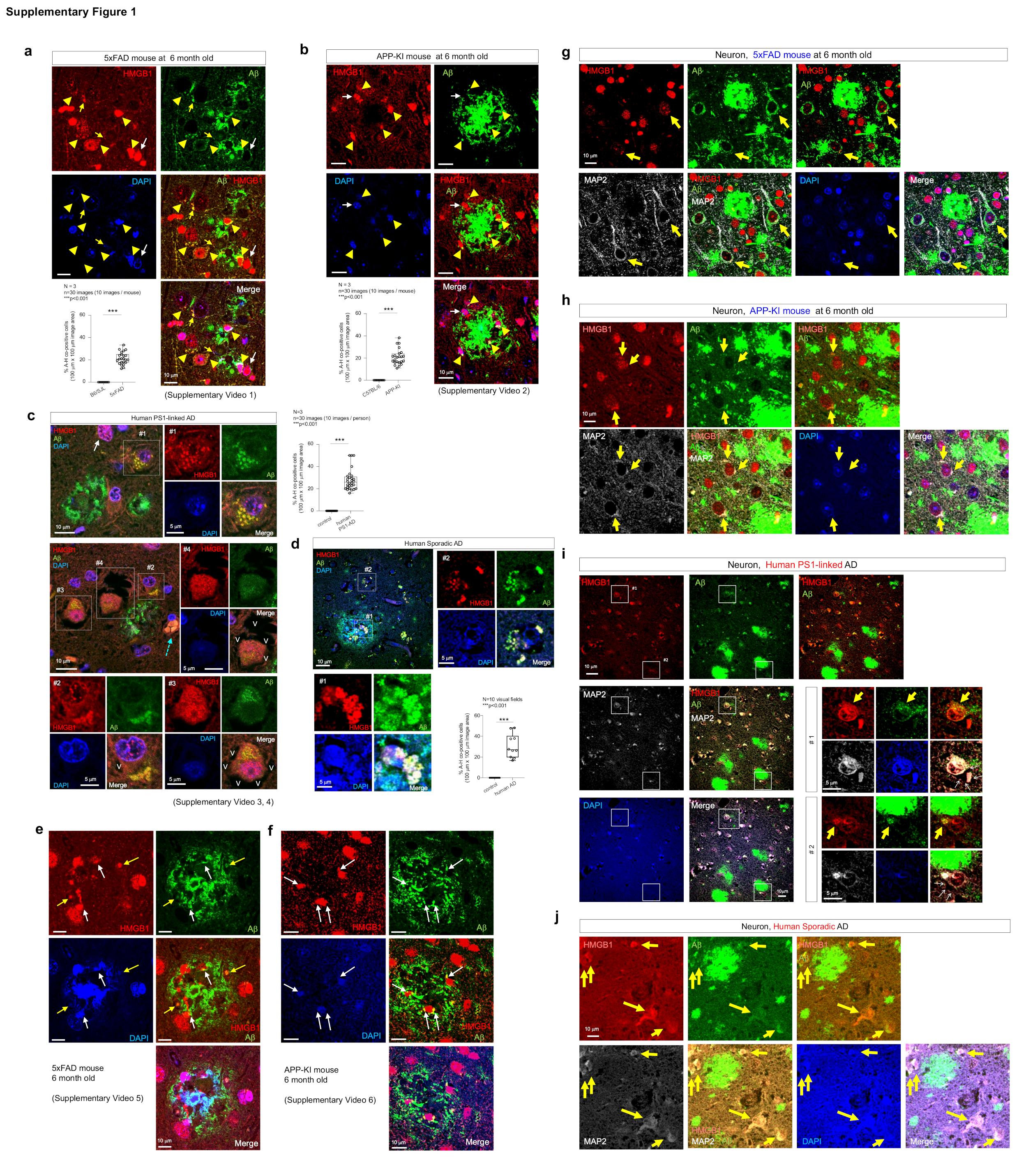

### Supplementary Figure 2

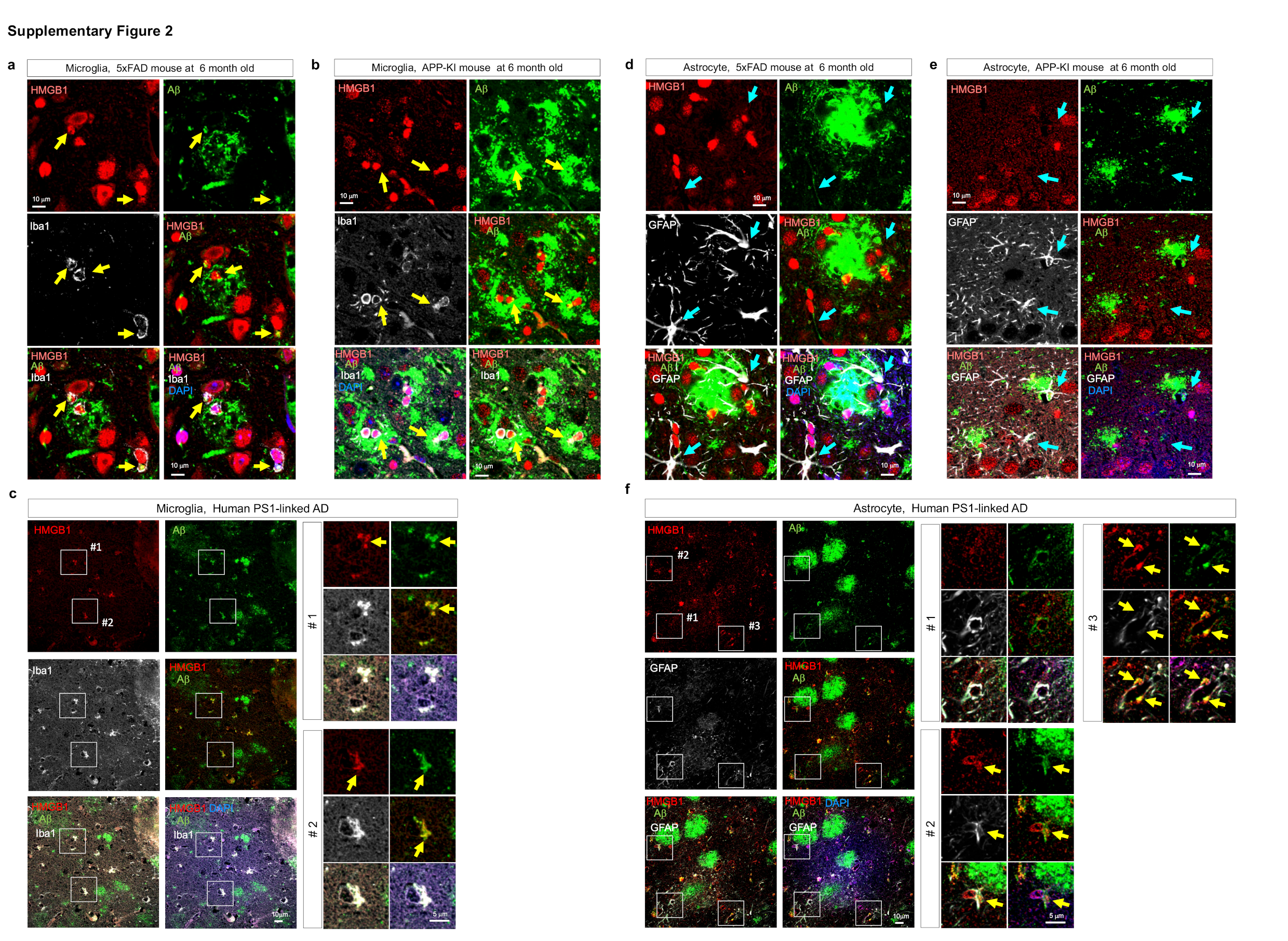

### Supplementary Figure 3

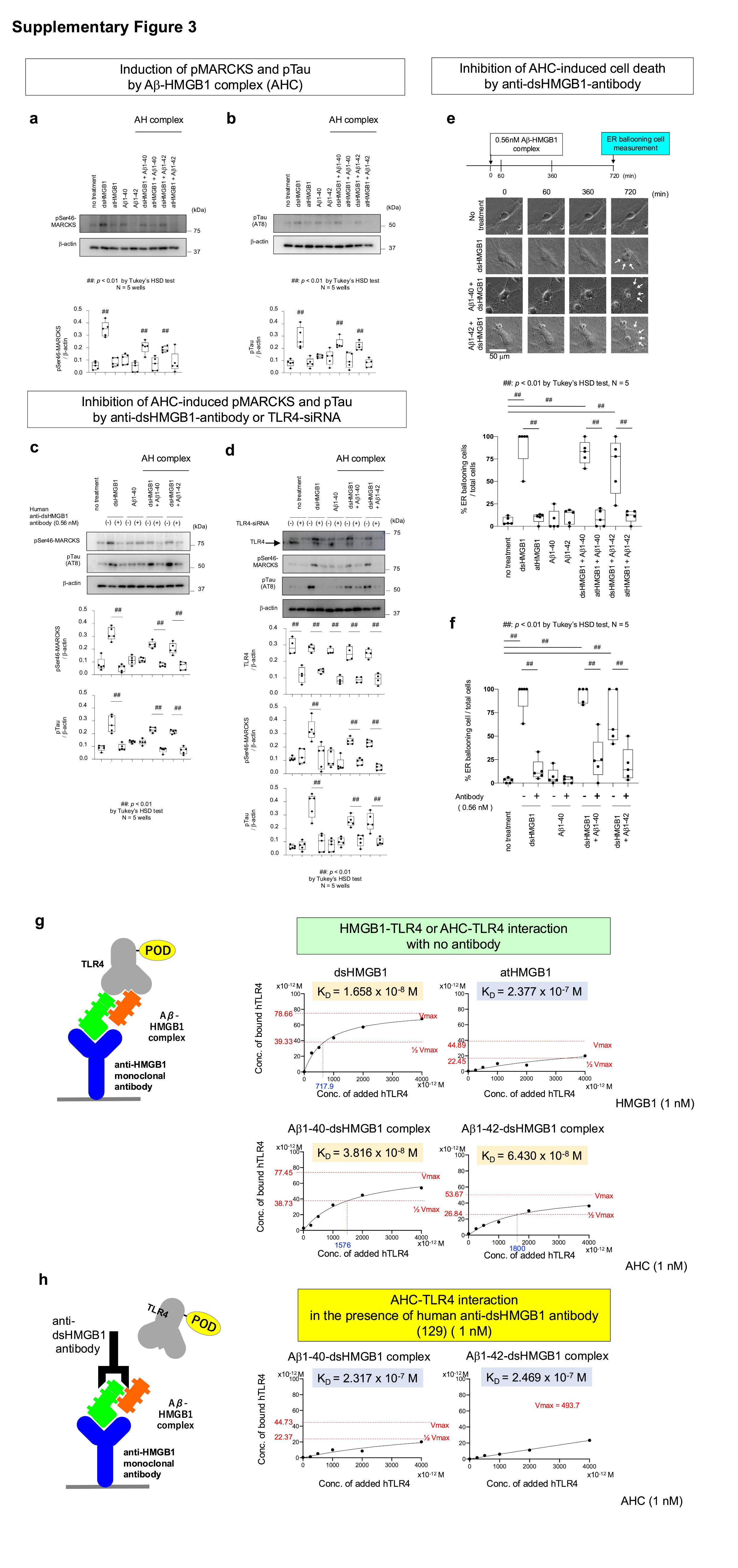

### Supplementary Figure 4

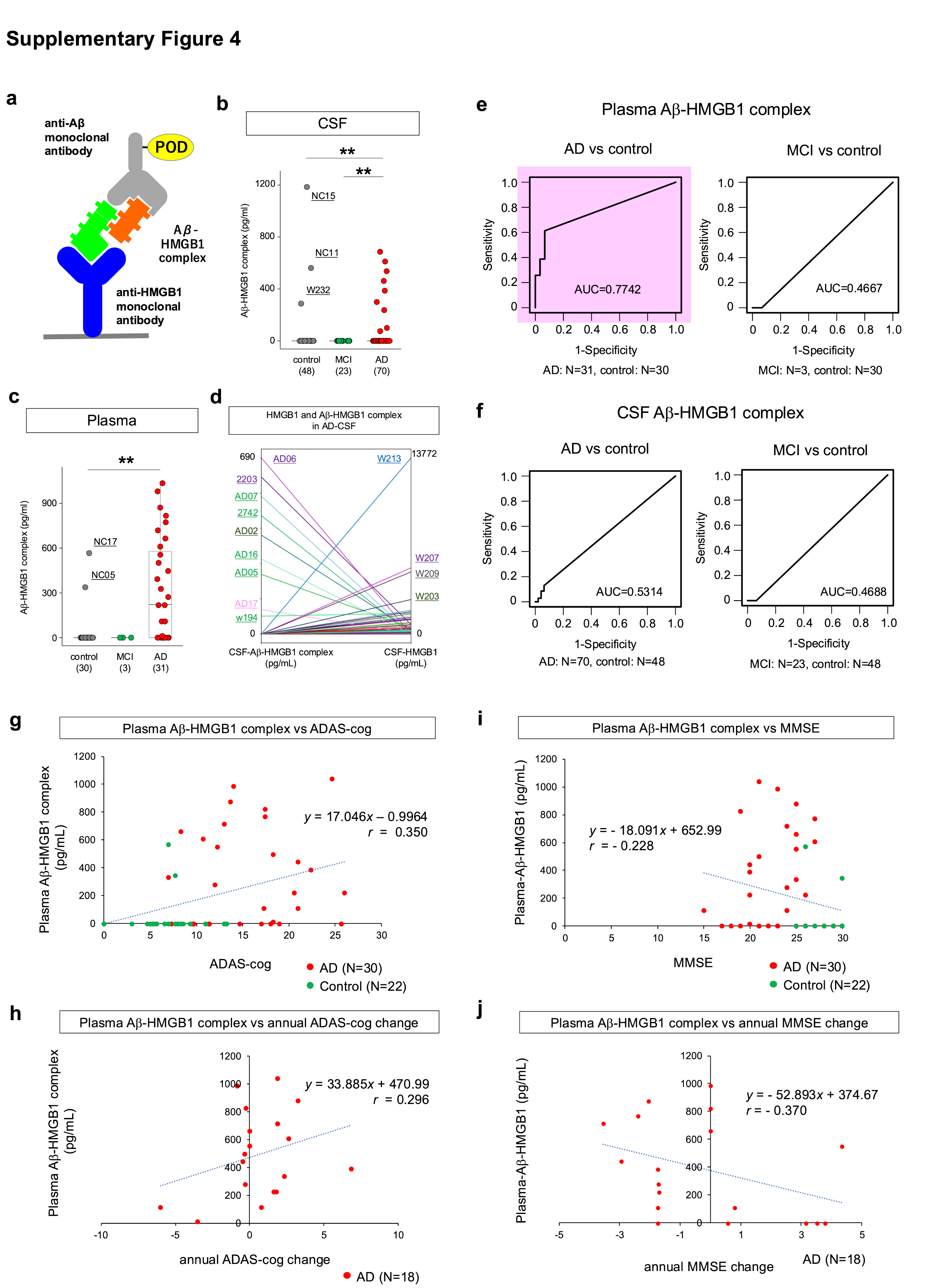

### Supplementary Figure 5

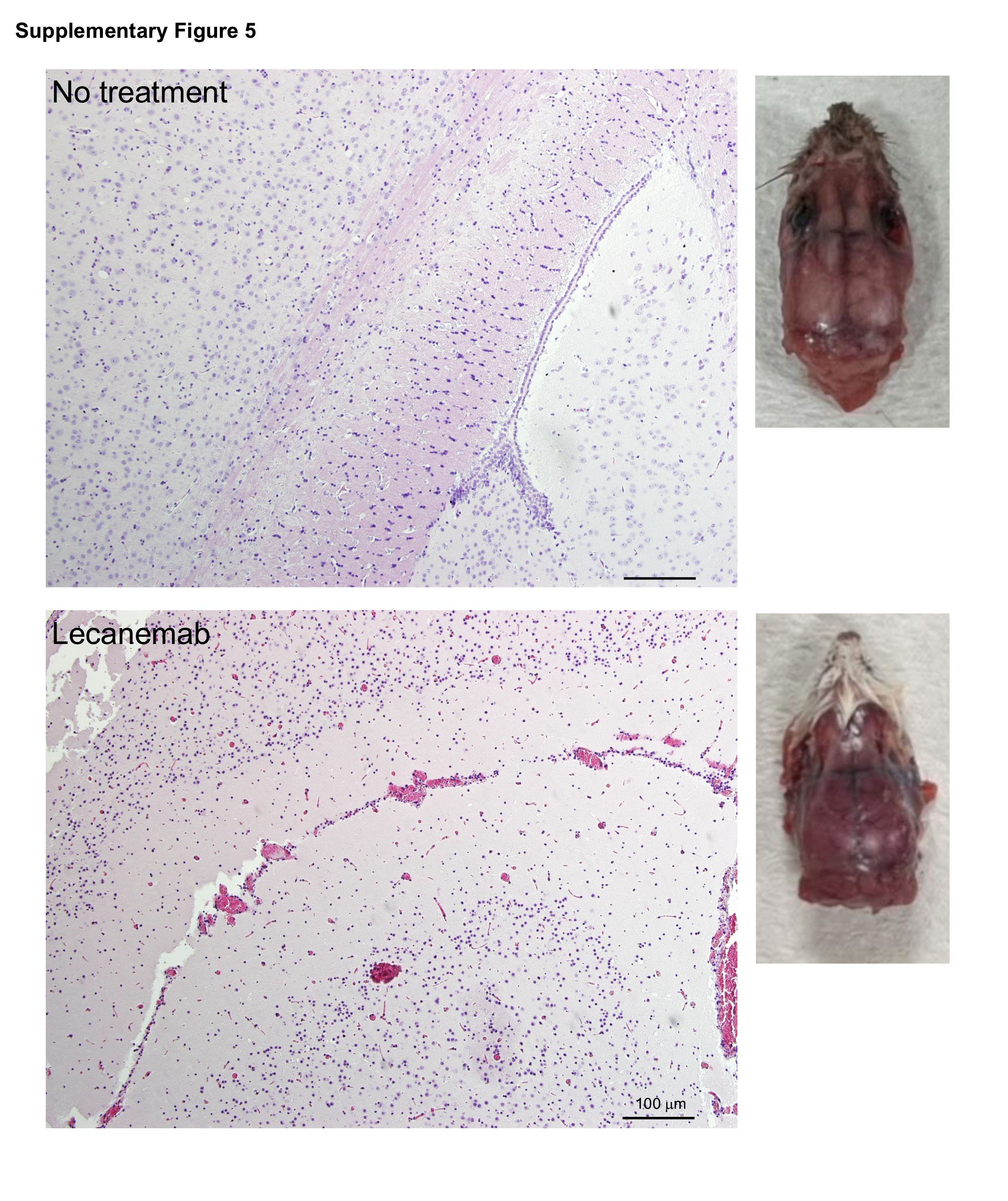

### Supplementary Figure 6

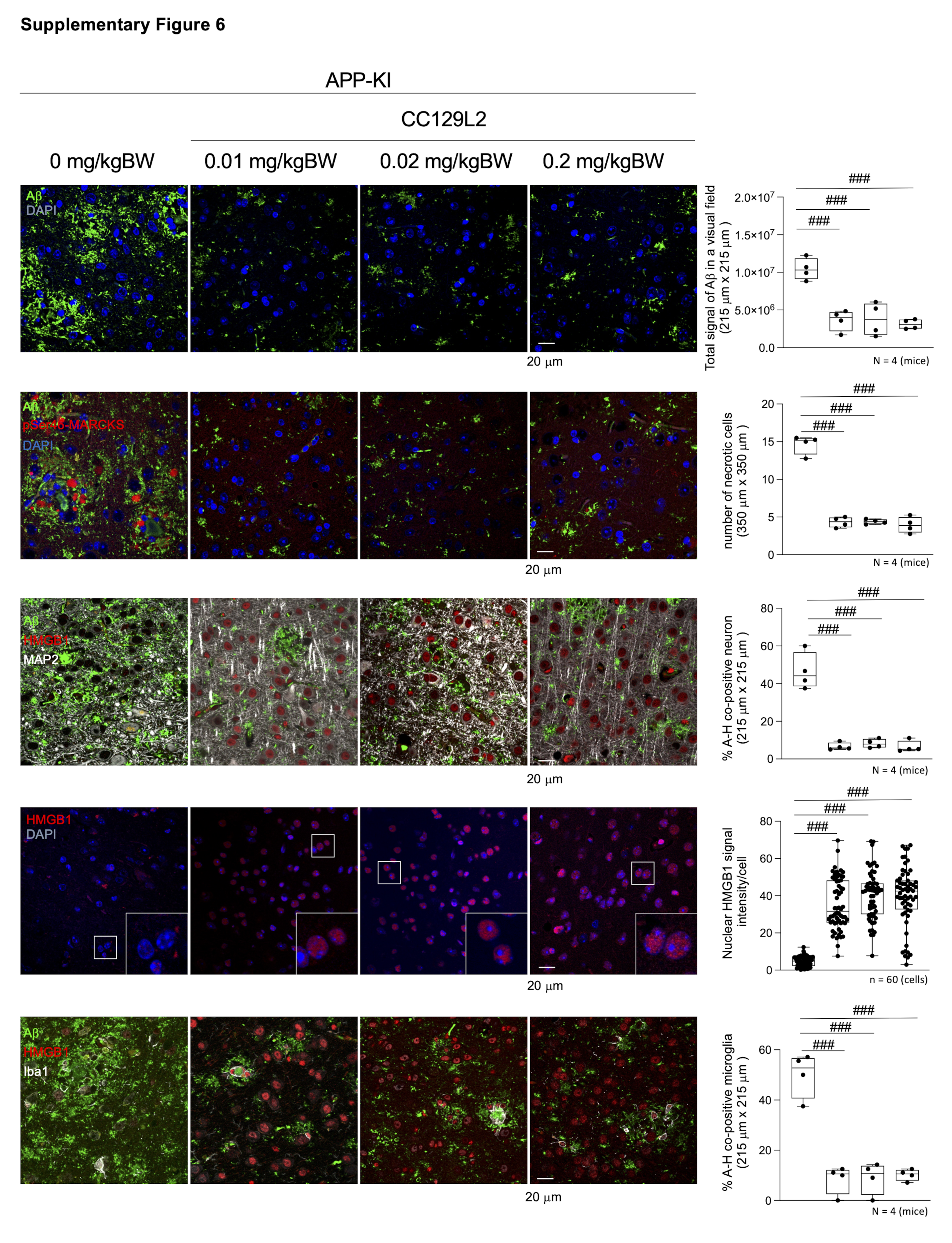

### Supplementary Figure 7

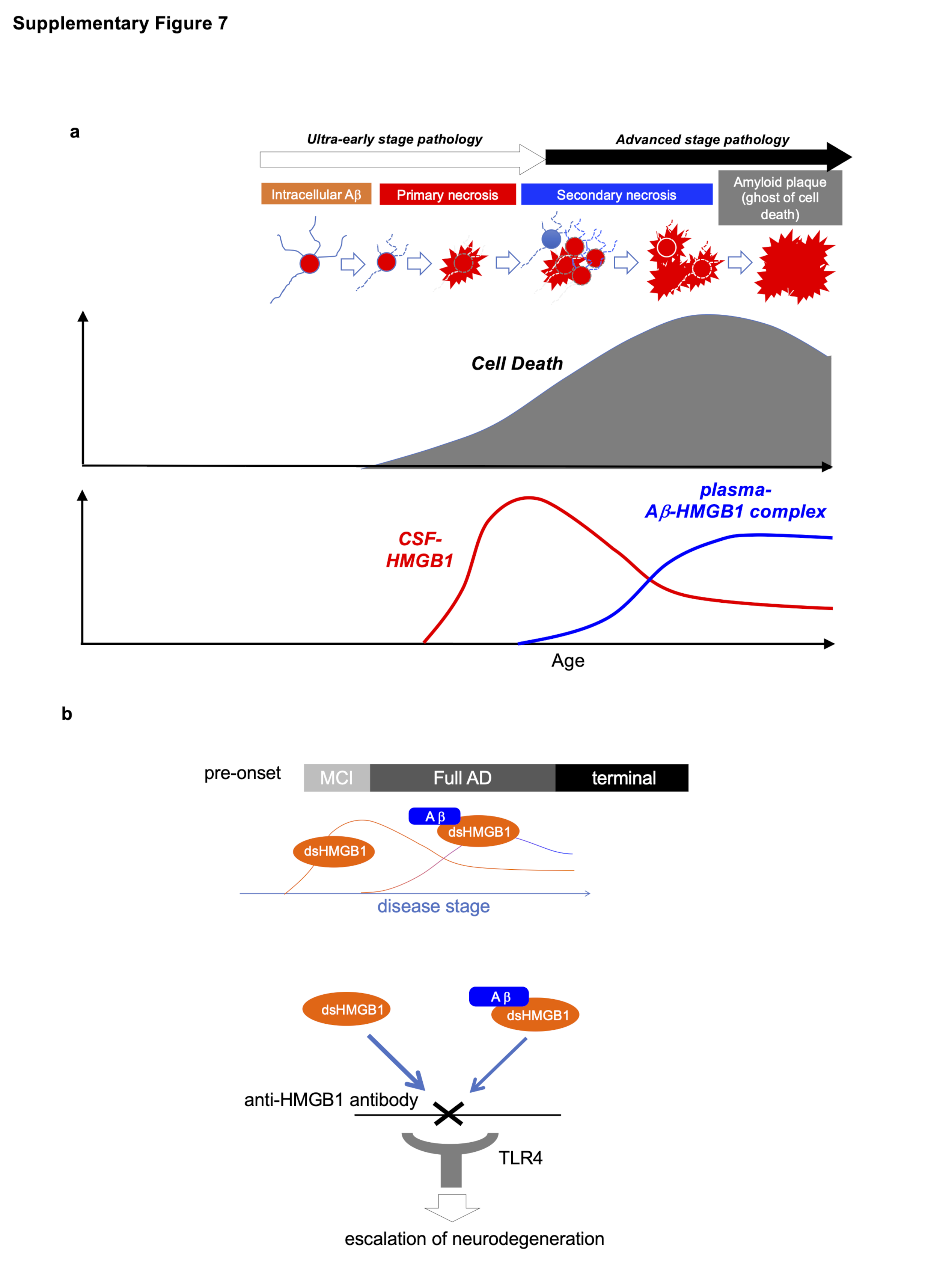

### Supplementary Table 1

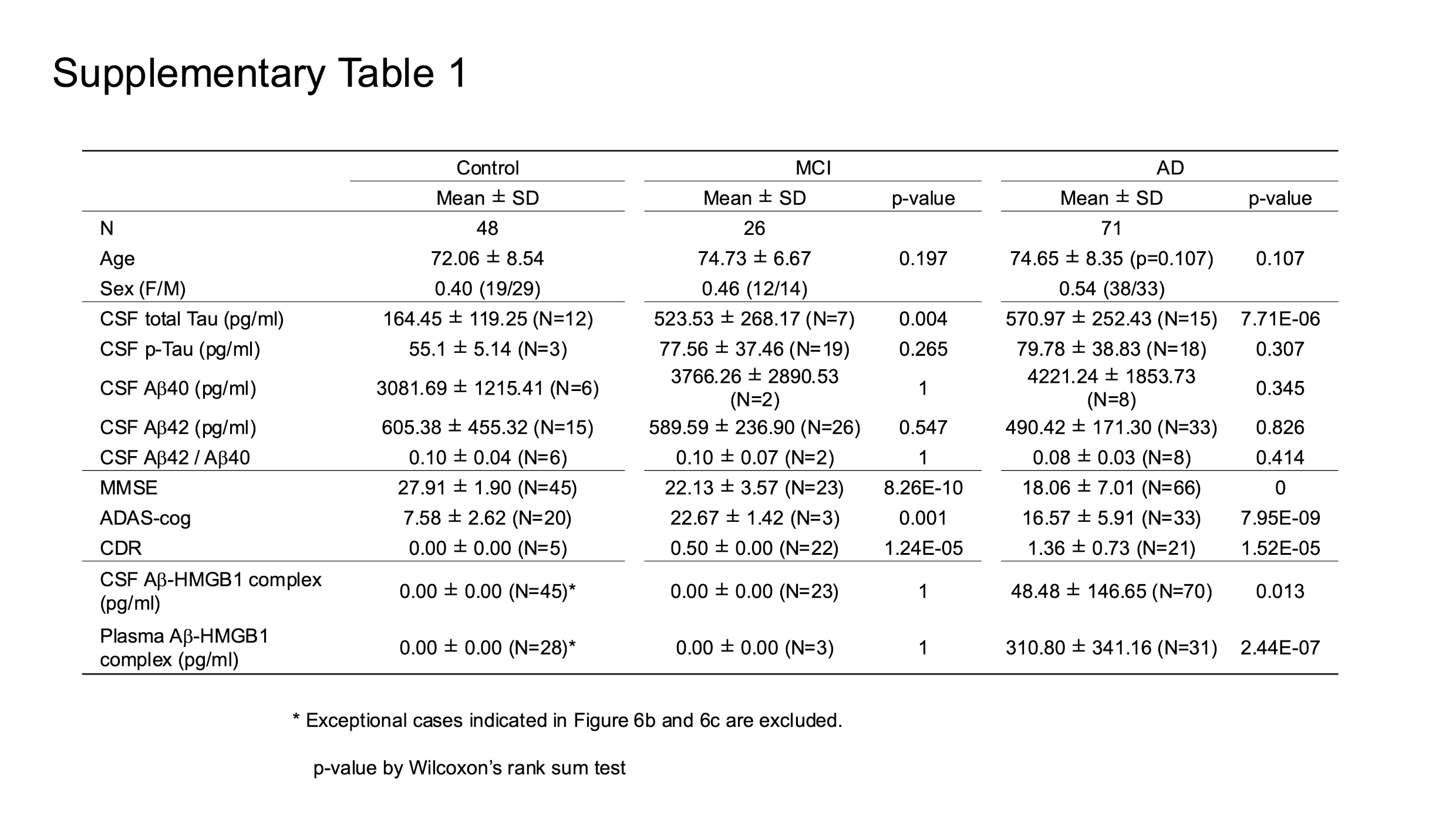

### Supplementary Table 2

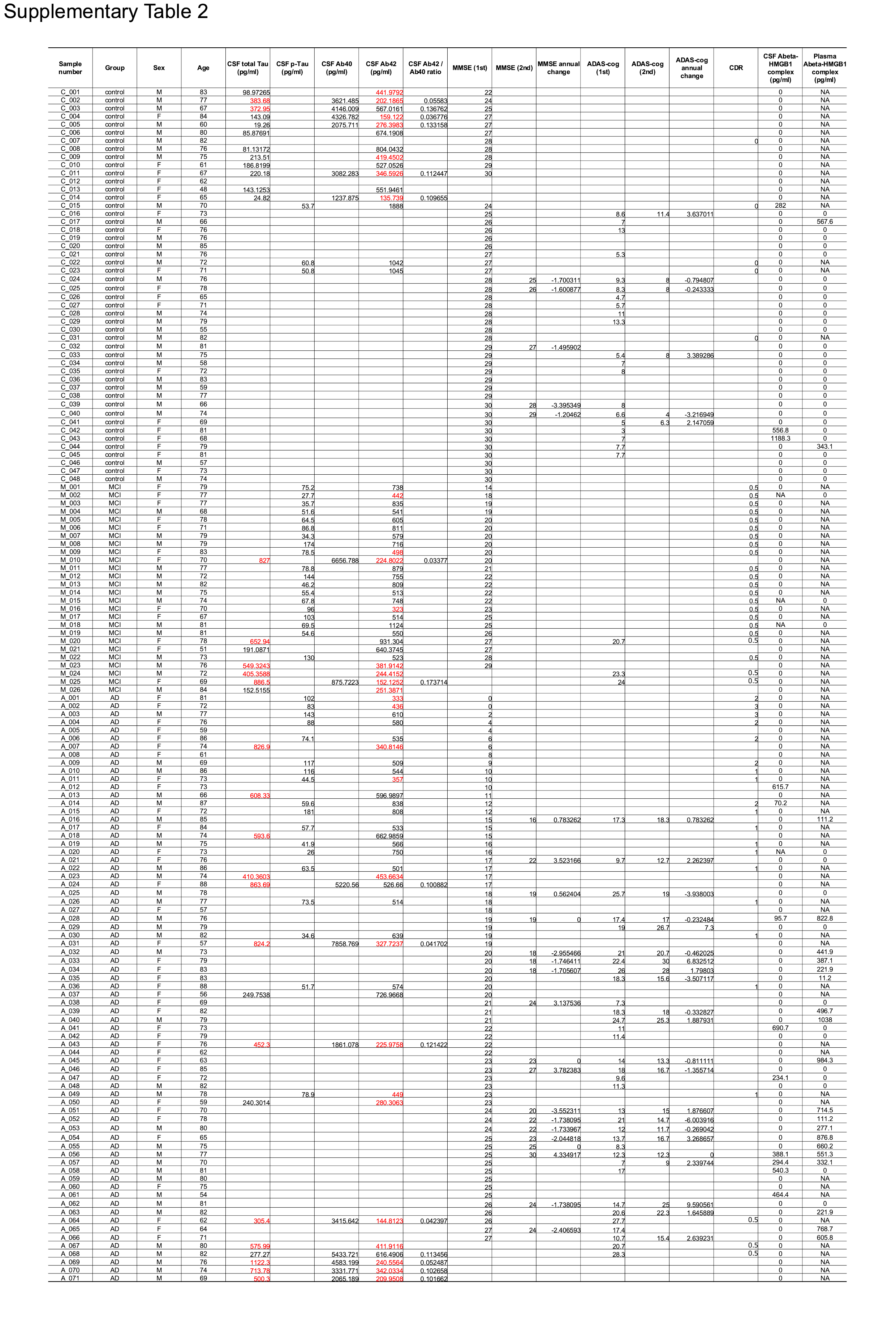
